# Computer code comprehension shares neural resources with formal logical inference in the fronto-parietal network

**DOI:** 10.1101/2020.05.24.096180

**Authors:** Y. Liu, J. Kim, C. Wilson, M. Bedny

**Affiliations:** Johns Hopkins University

## Abstract

Despite the importance of programming to modern society, the cognitive and neural bases of code comprehension are largely unknown. Programming languages might ‘recycle’ neurocognitive mechanisms originally used for natural languages. Alternatively, comprehension of code could depend on fronto-parietal networks shared with other culturally derived symbol systems, such as formal logic and math. Expert programmers (average 11 years of programming experience) performed code comprehension and memory control tasks while undergoing fMRI. The same participants also performed language, math, formal logic, and executive control localizer tasks. A left-lateralized fronto-parietal network was recruited for code comprehension. Patterns of activity within this network distinguish between “for” loops and “if” conditional code functions. Code comprehension overlapped extensively with neural basis of formal logic and to a lesser degree math. Overlap with simpler executive processes and language was low, but laterality of language and code covaried across individuals. Cultural symbol systems, including code, depend on a distinctive fronto-parietal cortical network.

## Introduction

In 1800, only twelve percent of the world’s population knew how to read, but today the world literacy rate is over eighty-five percent (https://ourworldindata.org/literacy). The ability to comprehend programming languages may follow a similar trajectory. Although only an estimated .5% of the world’s population is currently proficient at computer programming, the number of jobs that require programming continues to grow. Coding is essential in scientific fields and in areas as diverse as artistic design, finance, and healthcare. As many industries incorporate artificial intelligence or other information technologies, more people seek to acquire programming literacy. Yet the cognitive and neural mechanisms supporting coding remain largely unknown. Apart from its intrinsic and societal interest, programming is a case study of “neural recycling” (Dehaene & Cohen, 2007). Computer programming is a very recent cultural invention and the human brain is not evolutionarily adapted to support it. Studying the neural basis of code offers an opportunity to investigate how the human brain enables cultural inventions.

Hypotheses about how the human brain accommodates programming range widely. One recently popular view is that code comprehension recycles language processing mechanisms (Fedorenko, Ivanova, Dhamala, & Bers, 2019; Fitch, Hauser, & Chomsky, 2005; Pandža, 2016; Portnoff, 2018; Prat, Madhyastha, Mottarella, & Kuo, 2020). Computer languages borrow letters and words from natural language. In some programming languages, like Python, the meanings of the borrowed symbols (e.g., `if`, `return`, `print`) relate to the meanings of the same symbols in English. As in natural languages, the symbols of code combine generatively according to a set of rules (i.e., a formal grammar). Moreover, the grammars of language and that of code share common features, including recursive structure (Fitch et al., 2005). A recent study reported that individual differences in learning a second language predict aptitude in learning to program among novices (Prat et al., 2020). One possibility then is that coding recycles neurocognitive mechanisms involved in language processing (Peitek et al., 2018; Siegmund et al., 2014).

On the other hand, other culturally derived symbol systems, such as formal logic and math do not appear to depend on the same neural network as natural language. Like code, formal logic and mathematics borrow symbols from natural language and are also hierarchical and recursive (e.g. *(7*\**(7*\**(3*+*4*))). Unlike language, however, culturally derived symbol systems are explicitly taught later in life. Computer coding, mathematics and logic, also involve manipulation of arbitrary variables without inherent meaning (e.g. `X`, `Y`, `input`, `ii`) according to a set of learned rules (McCoy & Burton, 1988). Each symbol system, including code, has its own conventionalized variables and its own set of rules. In the case of code, the rules also differ somewhat across programming languages. The deductive rules of programming also share specific features with formal logic. For example, connectives (e.g., *“if*…*then”, “and”, “or”, “not”*) occur in both domains and have related meanings. Consider a function containing an `if` conditional written in Python,

~~~
def fun(input):
     result = “result: “
     if input[0]==“a”:
          result += input[0].upper()
return result
~~~

The value of the “`result`” variable depends on whether the “`input`” meets the specific conditions of the `if` statement. Similarly, in the logical statement “*If both X and Z then not Y*” the value of the result (*Y*) depends on the truth value of the condition “*both X and Z*”. One prediction, then, is that coding depends on similar neural resources to other culturally-derived symbol systems such as formal logic and math.

Rather than recruiting perisylvian fronto-temporal areas, mathematics and logic recruit a fronto-parietal network, including the dorsolateral prefrontal cortex and the intraparietal sulcus (IPS) as well as putative symbol representations (i.e. numberform area) in inferior temporal cortex (Amalric & Dehaene, 2016; Coetzee & Monti, 2018; Goel et al., 2007; Monti, Parsons, & Osherson, 2009). This fronto-parietal network overlaps partially with the so called central executive/working memory system, which is implicated in a variety of cognitive tasks that involve maintaining and manipulating information in working memory, processes that are part and parcel of understanding and writing code (Brooks, 1977; Duncan, 2010; Letovsky, 1987; Miller & Cohen, 2001; Soloway & Ehrlich, 1984; Weinberg, 1971; Zanto & Gazzaley, 2013)(for a review of the cognitive models of code comprehension, see (Von Mayrhauser & Vans, 1995)). The central executive system is usually studied using simple rule-based tasks, such as the multisource interference task (MSIT), Stroop and executive working memory (Banich et al., 2000; Bunge, Klingberg, Jacobsen, & Gabrieli, 2000; Bush & Shin, 2006; January, Trueswell, & Thompson-Schill, 2009; Milham et al., 2001; Woolgar, Thompson, Bor, & Duncan, 2011; Zanto & Gazzaley, 2013; Zhang, Kriegeskorte, Carlin, & Rowe, 2013). Logic and math activate a similar network but also have unique neural signatures. Within the prefrontal cortex, logic recruits more anterior regions associated with more advanced forms of reasoning and symbol manipulation (Coetzee & Monti, 2018; Ramnani & Owen, 2004).

The goal of the current study was to ask whether computer code comprehension has a consistent neural signature across people and if so whether this signature is similar to other culturally derived symbol systems (i.e., logic and math) or similar to natural language. Only a handful of studies have looked at the neural basis of coding (Duraes, Madeira, Castelhano, Duarte, & Branco, 2016; Floyd, Santander, & Weimer, 2017; Ikutani & Uwano, 2014; Peitek et al., 2018; Siegmund et al., 2014). Thus far results have been largely inconsistent, possibly due to the complexity of the tasks and absence of control conditions. Moreover, no prior study has directly compared the neural basis of code to other cognitive domains.

A group of expert programmers (average 11 years of programming experience) performed a code comprehension task while undergoing functional magnetic resonance imaging (fMRI). We chose a comprehension task rather than code writing or debugging partly because it could in principle be analogous to understanding language vignettes and because it is arguably simpler. On each *real code* trial, participants saw a short function definition, followed by an input and a possible output, and judged whether the output was valid. In *fake code* control trials, participants performed a memory task with arbitrary text. A fake function was generated by scrambling a real function per line at the level of word/symbol. Each fake function preserved the perceptual and lexical elements of a real function, but was devoid of syntactic structure. The *real code* condition contained two subtypes or ‘control structures’, `for` loops and `if` conditionals. We used multi-voxel-pattern analysis to decode `for` from `if` functions in order to test whether the code-responsive cortical system encodes code-relevant information. Finally, we examined the overlap of code comprehension with language (sentence comprehension), formal logic and mathematical tasks. We also tested overlap of code with the MSIT to determine whether the overlap with culturally derived symbol systems (i.e. logic and math) is more extensive than overlap with simpler experimentally defined rule-based tasks.

## Methods

### Participants

Seventeen individuals participated in the study, one did not complete the tasks due to claustrophobia, and another was excluded from analyses due to excessive movement (> 2mm). We report data from the remaining fifteen individuals (3 women, age range 20-38, mean age = 27.4, SD = 5.0). All participants had normal or corrected to normal vision, and no known cognitive or neurological disabilities. Participants gave informed consent according to procedures approved by the Johns Hopkins University Institutional Review Board.

All participants had at least 5 years of programming experience (range: 5-22, mean=10.7, SD=5.2), and at least 3 years of experience working with the programming language Python (range: 3-9, mean=5.7, SD=1.8).

### Behavioral pre-test

In addition to self-reported programming experience, Python expertise was evaluated with two out-of-the-scanner Python exercises (one easier and one more difficult) the week prior to the fMRI experiment. These exercises also served to familiarize participants with the particular Python expressions that would be used during the fMRI experiment.

The easier exercise consisted of three phases: 1. test, 2. recap and 3. re-test. During the first phase of the exercise (test), we evaluated participants’ knowledge of every built-in Python function that would occur during the fMRI experiment. Participants were asked to type the output of a single line of Python `print()` statement (e.g., for “`print(“3.14”.split(“1”))`” one should type “`[‘3.’, ‘4’]`”). On average participants answered M=82.9% (SD=6.9%) of the questions correctly (range: 70% - 96%). Since even expert programmers may not have used a particular function in the recent past, the “recap” phase of the exercise explicitly reviewed the definitions and purposes of all of the functions and expressions that would be used during the fMRI experiment. During the final re-test phase, participants were once again asked to type the output of a single line of Python for each function (M=92.0% (SD=7.5%), range: 72.4% - 100%).

The more difficult exercise evaluated the participants’ knowledge about when and how to use Python functions and expressions. Each participant answered sixteen questions, each consisting of a code snippet with a blank. A prompt was presented alongside the code snippet to explain what the snippet should output if executed. The participant was asked to fill in the blank in order to complete the code (see the sub-section “an example of the difficult out-of-the-scanner exercise” in the supplementary method). The questions were designed by the experimenter to cover some of the objectives specified in the exam syllabus of the Certified Associate in Python Programming Certification held by the Python Institute (https://pythoninstitute.org/certification/pcap-certification-associate/pcap-exam-syllabus/). On average, the participants got 64.6% (SD=16.6%) of the questions correct (range: 37.5% - 93.75%).

### fMRI task design and stimuli

#### Code Comprehension Experiment

In the *real code* comprehension trials, participants were presented with user-defined Python functions. Python comprehension was compared to a *fake code* control trials that consisted of incomprehensible scrambled Python functions (described in further detail below). To help participants distinguish between *real* and *fake code* trials, *real code* appeared in white text and *fake code* in yellow text.

Each trial consisted of three phases: *function* (24 seconds), *input* (6 seconds), and *question* (6 seconds) (Figure 1). First, participants viewed a Python function for 24 seconds, followed by a .5 second fixation-cross delay. During the *input* phase, the original code function re-appeared on the screen with a potential input below consisting of a single line character string (6 seconds). Participants were instructed to use the input to mentally derive the output of the code function during the *input* phase. After the *input* phase there was a .5 second fixation-cross delay followed by a proposed output along with the prompt “*TRUE?*” Participants were asked to determine whether the output was correct within 6 seconds. All trial phases had a shortening bar at the bottom of the screen indicating the remaining time during that particular phase of the trial. Each trial was followed by a 5-second inter-trial interval during which the text “*Your response is recorded. Please wait for the next trial*” was shown on the screen.

**Figure 1.**
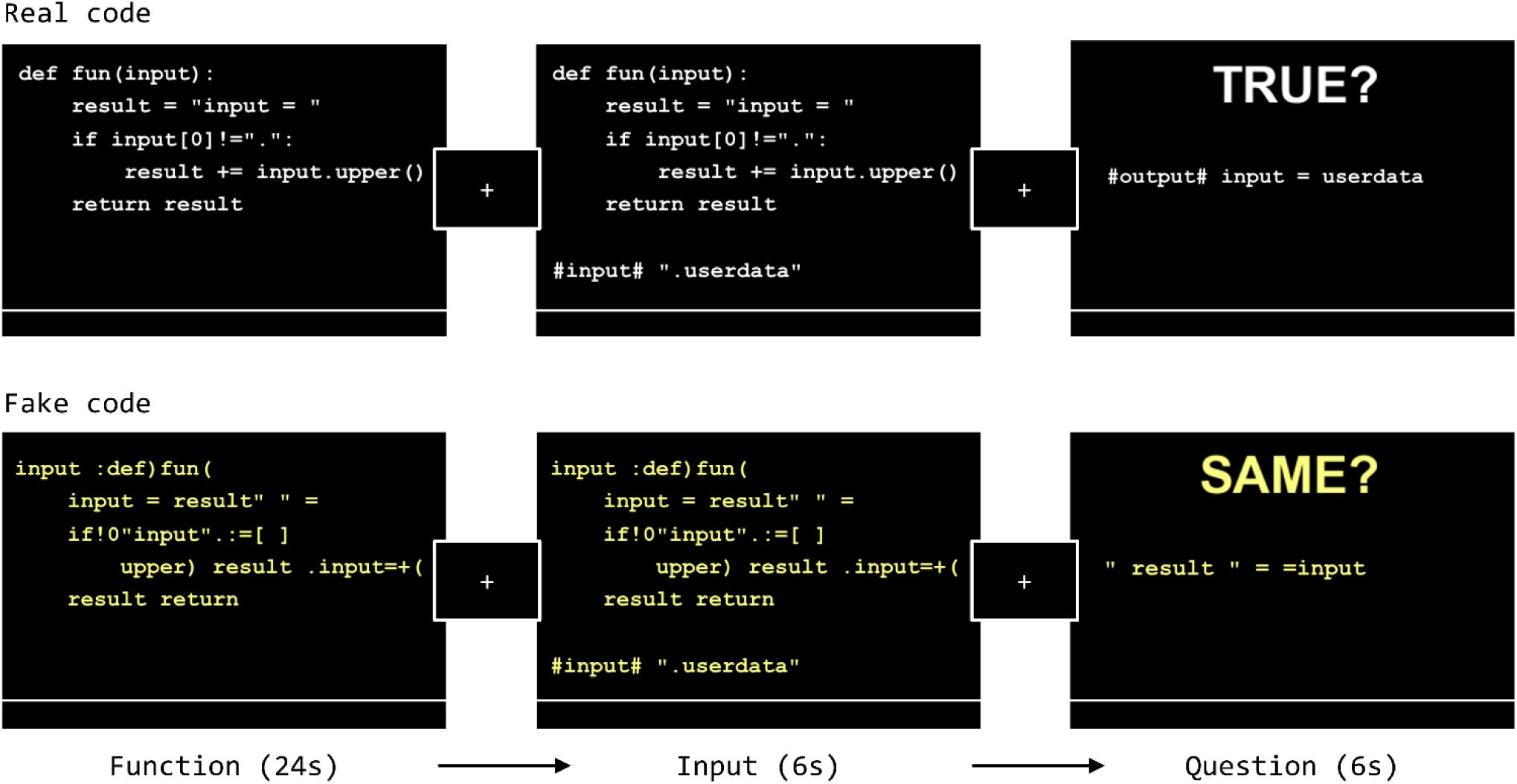
The experiment design. The FAKE function (bottom row) in this figure is created by scrambling the words and symbols in each line of the REAL function (top row). Note that for the purpose of illustration, the relative font size of the text in each screen shown in this figure is larger than what the participants saw during the actual MRI scan.

Each *real code* function consisted of five lines. The first line (`def fun(input):`) and the last (`return result`) were always the same. The second line always initialized the result variable, and the third and fourth lines formed a control structure (either a `for` loop or an `if` conditional) that may modify the value of the result. *Real code* trials were divided into two sub-conditions, `for` and `if`, according to the control structures the functions contained. Each condition included two variants of the `for` or `if` functions (see the sub-sections “detailed information about the stimuli” and “the two variants of each control structure” in the supplementary method). All functions took a letter string as input and performed string manipulation.

*Fake code* trials were analogous to the *real code* trials in temporal structure (i.e. function, input, question). However, no real code was presented. Instead, participants viewed scrambled text and were asked to remember it. During the *function* phase of a *fake code* trial, participants saw a scrambled version of a Python function. Scrambling was done within line at word and symbol level (Figure 1, bottom row). Because fake functions were derived from real functions, the words, digits and operators that existed in real functions were preserved but none of the lines comprised an executable Python statement. During the *input* phase, an additional fake input line appeared below the fake function. The fake input line didn’t interact with the fake function, the participants only had to memorize this line. During the *question* phase, a new character line appeared along with the prompt “*SAME?*” Participants judged whether this line had been presented during the *function* and *input* phases (including the additional input line), or it came from a different fake function. The answer was “true” on half of the *real* and *fake code* trials.

There were 6 task runs, each consisting of 20 trials, 8 *real* `if` code, 8 *real* `for` code and 4 *fake* code trials. Each participant saw a total of 48 `for` functions (24 per variant), 48 `if` functions (24 per variant), and 24 *fake* functions. After each run of the task, participants saw their overall percent correct and average response time. Participants were divided into two groups such that the variants of the functions were balanced across groups, and the same participant never saw different variants of the same function. The order of the presentation of the functions was pseudo-randomized and balanced across participants. In total, 192 *real* functions (96 per group) and 48 *fake* functions (24 per group) were used in the experiment. All the functions are listed in supplementary Table S1. We permuted the order of the functions systematically such that each participant saw a unique order (see the sub-section “the algorithm of the permutation of stimuli” in the supplementary method).

#### Localizer Tasks

During a separate MRI session, participants took part in two localizer experiments. A single experiment was used to localize responses to language, math and formal logic using each condition as the control for the others: language/math/logic localizer. The task design was adapted from Monti et al. 2009, 2012 (Kanjlia, Lane, Feigenson, & Bedny, 2016; Monti et al., 2009; Monti, Parsons, & Osherson, 2012). On language trials, participant judged whether two visually presented sentences, one in active and one in passive voice, had the same meaning (e.g. “*The child that the babysitter chased ate the apple*” vs “*The apple was eaten by the babysitter that the child chased*”). On math trials, participant judged whether the variable X had the same value across two equations (e.g. “*X minus twenty-five equals forty-one*” vs “*X minus fifty-four equals twelve*”). On formal logic trials, participant judged whether two logical statements were consistent, where one statement being true implied the other also being true (e.g. “*If either not Z or not Y then X*” vs “*If not X then both Z and Y*”).

Each trial began with a 1-second fixation cross. One pair member appeared first, the other 3 seconds later. Both statements remained on screen for 16 seconds. Participants pressed the right or left button to indicate true/false. The experiment consisted of 6 runs, each containing 8 trials of each type (language/math/logic) and 6 rest periods, lasting 5 seconds each. All 48 statement pairs from each condition were unique and appeared once throughout the experiment. In half of the trials, the correct answer was “true”. Order of trials was counterbalanced in two lists across participants.

Note that although all of the tasks in the language/math/logic localizer contain language stimuli, previous studies show that sentences with content words lead to larger responses in the perisylvian fronto-temporal language network than spoken equations or logical statements with variables (Kanjlia et al., 2016; Monti et al., 2009, 2012). The perisylvian fronto-temporal language network shows enhanced activity for stimuli that contain meaningful lexical items *and* sentence level syntax (e.g., (Fedorenko et al., 2016)). Furthermore, previous studies found that responses to language, logic and math when compared to each other were similar to what was observed for each domain relative to independent control conditions (e.g. sentences relative to lists of non-words for language, and hard vs. easy logic problems) (Kanjlia et al., 2016; Monti et al., 2009, 2012).

The multi-source interference task (MSIT) was adapted from Bush and Shin (Bush & Shin, 2006) to engage executive control processes and localize the multiple demand network. On each trial, a triplet of digits was shown on the screen, and two of the digits were the same. The participant pressed buttons (1, 2, or 3) to indicate the identity of the target digit which is different from the two distractors. For example, for “131” the correct response is “3”; for “233” it is “2”. The participants always pressed button “1”, “2”, and “3” with their index, middle, and ring fingers, respectively.

MSIT consisted of “control” blocks and “interference” blocks, each containing 24 trials (1.75 seconds each). On interference trials, the location of the target digit was never consistent with the identity of the digit (e.g., trials such as “133” or “121” never existed), thus giving rise to an interference. On control trials, the distractors were always “0”, and the target digit was always at the same location as its identity. In other words, there were only three kinds of control trials, “100”, “020”, and “003”.

The participant performed 2 runs of MSIT. Each run began with 15 seconds of fixation, followed by 4 control blocks and 4 interference blocks interleaved, and ended with another 15 seconds of fixation. Both an interference block and a control block lasted for 42 seconds. The order of the blocks was balanced both within and between participants. The order of the trials were arranged such that all 12 interference trials appeared exactly twice in an interference block, and all 3 control trials appeared exactly 6 times in a control block. Two same trials never came in succession, and the order of the trials was different across all 8 blocks of the same kind.

### Data acquisition

MRI data were acquired at the F.M. Kirby Research Center of Functional Brain Imaging on a 3T Phillips Achieva Multix X-Series scanner. T1-weighted structural images were collected in 150 axial slices with 1mm isotropic voxels using the magnetization-prepared rapid gradient-echo (MP-RAGE) sequence. T2*-weighted functional BOLD scans were collected in 36 axial slices (2.4 x 2.4 x 3mm voxels, TR = 2s). We acquired the data in one code comprehension session (6 runs) and one localizer session (2 runs of MSIT followed by 6 runs of language/matth/local judgment), with the acquisition parameters being identical for both sessions.

The stimuli in both the code comprehension and localizer sessions were presented using stand-alone PsychoPy3 software (https://www.psychopy.org/). The stimuli were projected to a rear projection screen cut to fit the scanner bore with an Epson PowerLite 7350 projector. The resolution of the projected image was 1600 x 1200. The participant viewed the screen via a front-silvered, 45∘inclined mirror attached to the top of the head coil.

### fMRI data preprocessing and general linear model (GLM) analysis

Data were analyzed using Freesurfer, FSL, HCP workbench, and custom in-house software written in Python. Functional data were motion corrected, high-pass filtered (128 s), mapped to the cortical surface using Freesurfer, spatially smoothed on the surface (6-mm FWHM Gaussian kernel), and prewhitened to remove temporal autocorrelation. Covariates of no interest were included to account for confounds related to white matter, cerebral spinal fluid, and motion spikes.

The four *real* function code (`for`1, `for`2, `if`1, `if`2) and *fake code* conditions were entered as separate predictors in a GLM after convolving with a canonical hemodynamic response function and its first temporal derivative. Only the images acquired during the twenty-four-second *function* phase were modeled.

For the localizer experiment, a separate predictor was included for each of the three conditions (language, math, and logic) modeling the 16 seconds during which the statement pair was presented, as well as a rest period (5 seconds) predictor. In the MSIT task, the interference condition and the control condition were entered as separate predictors.

Each run was modeled separately, and runs were combined within each subject using a fixed-effects model (Dale, Fischl, & Sereno, 1999; Smith et al., 2004). For the group-level analysis across participants, random-effects models were applied, and the models were corrected for multiple comparisons at vertex level with p < 0.05 false discovery rate (FDR) across the whole brain. A nonparametric permutation test was further implemented to cluster-correct at p < 0.01 family-wise error rate.

### ROI definition

For each participant, 4 code-responsive functional ROIs were defined to be used in the MVPA analysis. First, random-effects whole-brain univariate analysis for the *real* > *fake code* contrast revealed 4 major clusters in the left hemisphere: the intraparietal sulcus (IPS), the posterior middle temporal gyrus (pMTG), the lateral prefrontal cortex (PFC), and the early visual cortex (Occ). These clusters were used to define group search spaces. Each search space was defined by combining parcels from Schaefer et al. that encompassed each cluster (400-parcel map, (Schaefer et al., 2018)). Next, individual functional ROIs were defined within these clusters by taking the top 500 active vertices for the *real* > *fake* contrast within each participant.

### MVPA

MVPA was used to distinguish `for` and `if` functions based on the spatial activation pattern in code-responsive ROIs. Specifically, we used the support vector machine (SVM) implemented in the Python toolbox Scikit-learn (Chang & Lin, 2011; Pedregosa et al., 2011).

For each participant, the spatial activation pattern for each function was defined as the beta parameter estimation of a GLM with each function entered as a separate predictor. Within each ROI in each participant, the 96 spatial patterns elicited by the real functions were collected. Normalization was carried out separately for `for` condition and `if` condition such that in either condition, across all vertices and all functions, the mean is set to 0 and standard deviation to 1. The purpose of the normalization is to eliminate the difference in the baselines in both conditions while preserving the spatial patterns.

The whole dataset was split into a training test (90%, 86 functions) and a testing set (10%, 10 functions), where in each set, half of the patterns came from `for` functions. A linear SVM (regularization parameter C = 5.0) was trained on the training set and tested on the testing set. Classification was carried out on 100 different train-test splits, and the average accuracy value was recorded as the observed accuracy.

We tested the classifier performance against chance (50%) using a combined permutation and bootstrapping approach (Schreiber & Krekelberg, 2013; Stelzer, Chen, & Turner, 2013). We derived the t-statistic of the Fisher-z transformed accuracy values against chance (also Fisher-z transformed). The null distribution for each participant was generated by first shuffling the condition labels 1,000 times, then computing the mean accuracy derived from the 100 train-test split of each shuffled dataset. Then, a bootstrapping method was used to generate an empirical distribution of the t-statistics. In each of the 10^6^ iterations of the bootstrapping phase, one Fisher-z transformed null accuracy value (out of 1,000) per participant was randomly selected, and a one sample t-test was applied to the null sample. The empirical p-value of the real t-statistic was defined as the proportion of the null t-statistics greater than the real value.

### Overlap analysis

For each participant, in each hemisphere, we used cosine similarity to quantify the overlap of the activated vertices between code comprehension and each of the four localizer contrasts: language (language > math), math (math > language), logic (logic > language), and multi-source interference (hard > easy). First, we generated the binary activation map for each contrast. A vertex was assigned the value 1 if the significance of its activation is above the 0.05 (FDR corrected) threshold, and 0 otherwise. Each binary map was regarded as a vector, and the cosine similarity between two vectors (e.g., code comprehension and logic) was defined as the inner product of the vectors divided by the product of their respective lengths (norms). The cosine similarities of code to each of the localizer tasks was then compared using repeated-measure ANOVA and post-hoc pairwise comparisons with false discovery rate (FDR) correction.

The empirical lower bound was calculated separately for each localizer task to account for differences in the number of activated vertices across tasks. For each participant, for each localizer task, we computed the cosine similarity between the binary map for code comprehension and a shuffled binary map for each localizer task. This step was repeated 100 times to generate the null distribution of the similarity values.

We used a bootstrapping approach to test whether each observed cosine similarity value was significantly above the empirical lower bound. For each localizer task, we randomly selected one similarity value from the null distribution of one participant and computed a null group mean similarity. This step was repeated 10^6^ times to derive the null distribution of the null group mean similarity. The empirical p-value of the real group mean similarity was defined as the proportion of the null values greater than the real value.

We operationalized the empirical upper bound as the cosine similarity of code comprehension and itself. For each participant, we split the data for code comprehension in half, ran a GLM for each half, and derived two binary maps whose cosine similarity was computed. We averaged all the similarity values resulting from the 10 possible splits of the 6 runs and across all participants.

## Results

### Behavioral results

Accuracy was similar across *real* and *fake code* trials (*real* M=92%, SD=0.045; *fake* M=0.90, SD=0.069; binary logistic mixed regression, *real* to *fake* odds ratio β = 1.27; Wald’s z statistic, z = 1.21; p = 0.23). Accuracy was also similar across `if` and `for` trials (`if` M=0.92, SD=0.076; `for` M=0.92, SD=0.056; `if` to `for` odds ratio β = 0.95; Wald’s z statistic, z = -0.28; p = 0.77). Participants were slower to respond to *fake* as compared to *real code* trials (*real* M=1.73 sec, SD=0.416; *fake* M=2.03 sec, SD=0.37; t(73) = 2.329, p = 0.023) and slower to respond to `for` as compared to `if` trials (`for` M=1.85 sec, SD=0.46; `if` M=1.60 sec, SD=0.44; t(58) = 2.127, p = 0.038) (Figure S1).

In the language/math/logic localizer task, participants performed least accurately on logic trials, followed by math and language (logic M=0.82, SD=0.13; math M=0.94, SD=0.028; language M=0.98, SD=0.023; one-way-ANOVA, F(2, 42) = 18.29, p < 0.001). Participants were slowest to respond to logic trials, followed by math trials, and fastest on the language trials (logic M=6.47 sec, SD=2.42; math M=4.93 sec, SD=1.32; language M=4.03, SD=1.27; one-way-ANOVA F(2, 42) = 7.42, p = 0.0017) (Figure S1).

In the MSIT experiment, hard and easy conditions did not differ in terms of accuracy (hard M=0.97, SD=0.038; easy M=0.98, SD=0.034; t(28) = -1.363, p = 0.18), but the hard trials took significantly longer to respond to than the easy trials (hard M=0.792 sec, SD=0.092; easy M=0.506 sec, SD=0.090; t(28)=8.59, p < 0.001) (Figure S1).

### fMRI results

#### Code comprehension experiment

As compared to *fake code, real code* elicited activation in a left-lateralized network of regions, including the lateral PFC (middle/inferior frontal gyri, inferior frontal sulcus; mainly BA 44 & 46, with partial activation in BA 6, 8, 9, 10, 47), the parietal cortex (the IPS, angular and supramarginal gyri; BA 7) and the pMTG and superior temporal sulcus (BA 22 & 37). Activity was also observed in early visual cortices (Occ) (p < 0.01 FWER, Figure 2) (supplementary Table S2).

**Figure 2.**
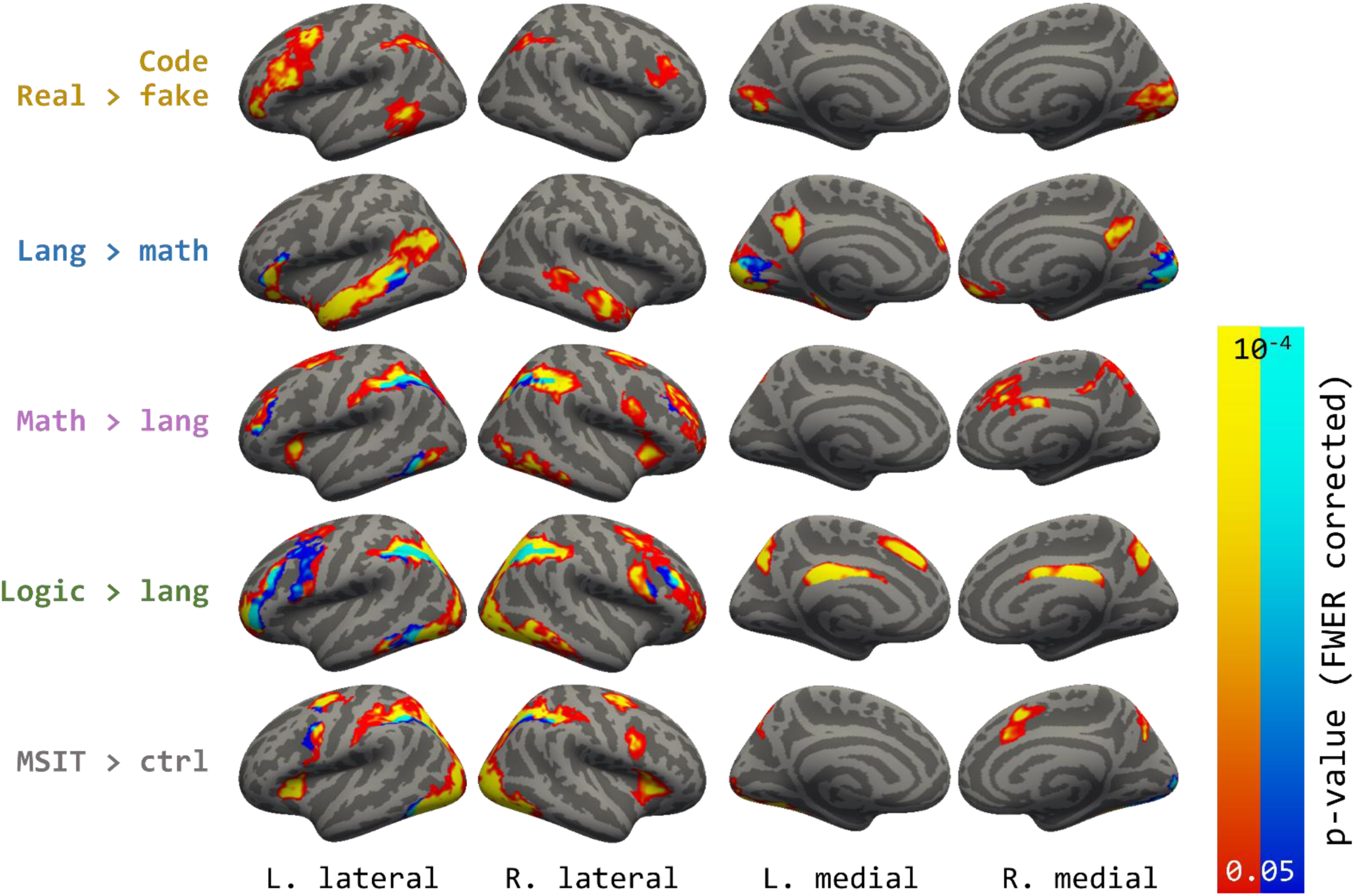
Whole brain contrasts. Areas shown are p<0.05 cluster-corrected p-values, with intensity (both warm and cold colors) representing uncorrected vertex-wise probability. In the maps for each localizer contrast, both warm and cold colors indicate activated vertices in the contrast, with the cold color labelling the overlap with the code contrast.

MVPA analysis revealed that `for` and `if` functions could be distinguished based on patterns of activity within PFC (accuracy = 64.7%, p < 0.001), IPS (accuracy = 67.4%, p < 0.001) and pMTG (accuracy = 68.4%, p < 0.001). `for` and `if` functions could also be distinguished within the early visual cortex (accuracy = 55.7%, p = 0.015), however, decoding accuracy was lower than in the other regions (F(3, 56) = 4.78, p = 0.0048) (Figure 3).

**Figure 3.**
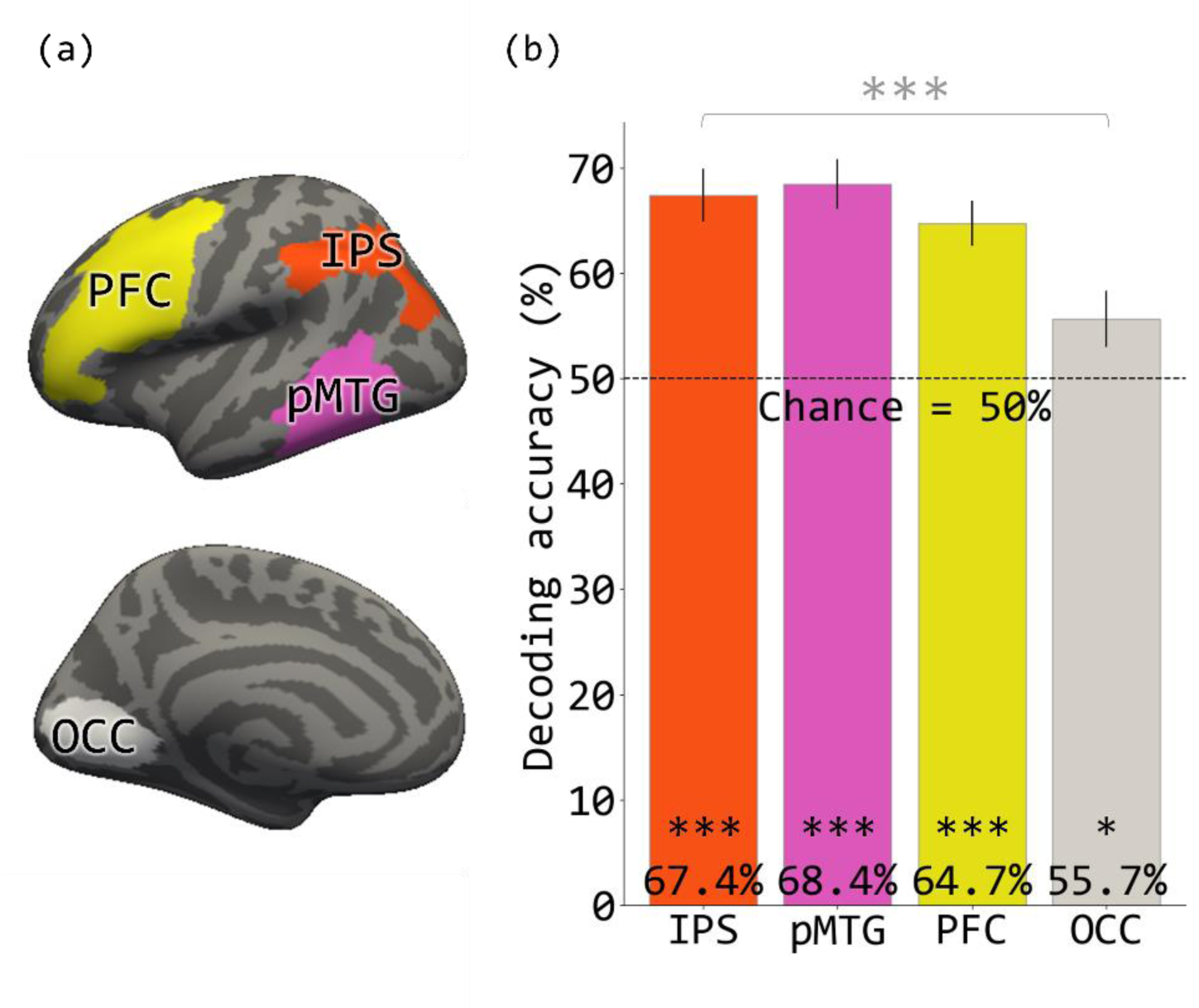
(a) The four search spaces (IPS, pMTG, PFC, OCC in the left hemisphere) within which functional ROIs were defined for the MVPA. (b) The MVPA decoding accuracy in the four ROIs. Error bars are mean ± SEM. *P<0.05. ***P<0.001.

#### Overlap between code comprehension and other cognitive domains

The language/math/logic localizer task activated previously identified networks involved in these respective domains. Responses to language were observed in a left perisylvian fronto-temporal language network, to math in parietal and anterior prefrontal areas as well as posterior aspect of the inferior temporal gyrus, and finally to logic, like math, in parietal and anterior prefrontal areas as well as posterior aspect of the inferior temporal gyrus. Logic activated more anterior and more extensive regions in prefrontal cortex than math. The MSIT hard > easy contrast also activated a fronto-parietal network including the IPS, however, the activation in the lateral frontal cortex was posterior and close to the precentral gyrus. (Figure 4, see supplementary Table S2 for full description of activation patterns associated with language, logic, math and MSIT). Note that although in the current experiment logic, math and language were compared to each other, the networks observed for each domain are similar to those previously identified with other control conditions (e.g. lists of non-words for language and hard vs. easy contrast in a logic task) (e.g.(Coetzee & Monti, 2018; Fedorenko, Behr, & Kanwisher, 2011)).

**Figure 4.**
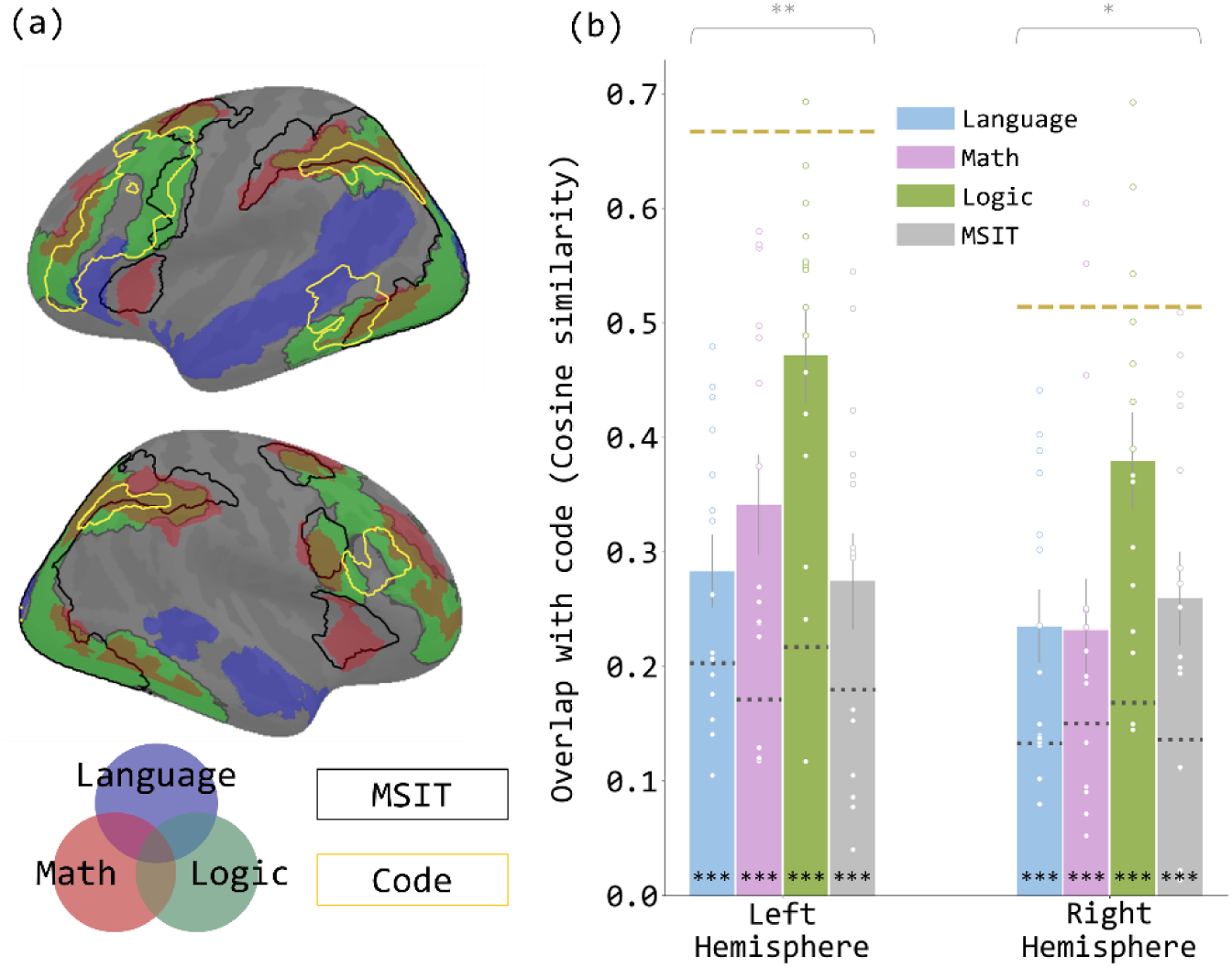
(a) Brain map with the activated regions in the five contrasts reported in Figure 2 overlain. The language network is shown in transparent blue, math in transparent red, and logic in transparent green. The regions activated in the MSIT contrast are enclosed in black outlines, and the code-responsive regions are enclosed in yellow outlines. (b) Cosine similarity between code contrast and each localizer contrast, in each hemisphere. Each dot represents the data from one participant. The dotted line on each bar indicates the null similarity between code contrast and the given localizer contrast. The yellow dashed line in each hemisphere indicates the empirical upper bound of the cosine similarity, the similarity between code comprehension and itself, averaged across participants. Error bars are mean ± SEM. *P<0.05. **P<0.01. ***P<0.001

Because code comprehension was highly left lateralized, overlap analyses focused on the left hemisphere. Right hemisphere results are reported in the sub-section “overlap analysis in the right hemisphere” in the supplementary results. Code comprehension (*real* > *fake*) overlapped significantly above chance with all localizer tasks: logic, math, language and MSIT (each task compared to chance p’s < 0.001, compared to code split-half overlap p’s < 0.005) (Figure 4). The degree of overlap differed significantly across tasks (repeated-measures ANOVA: F(3,42) = 5.04, p = 0.0045). Code comprehension overlapped most with logic (logic > language), followed by math and least with MSIT and language (Figure 4). Overlap with logic was significantly higher than with all other tasks, while the overlaps with the other three tasks (language, math, MSIT) were statistically indistinguishable from each other (post-hoc paired t-tests, FDR-corrected p’s < 0.05) (Supplementary Table S3).

The overlap of code with logic and math was observed in the IPS, PFC and a posterior portion of the inferior temporal gyrus (IT). PFC overlap was localized to the anterior middle frontal gyrus (aMFG, BA 46) and posteriorly in the precentral gyrus (BA 6). Overlap of code and the MSIT (hard > easy) was also observed in the IPS, precental gyrus and a small portion of the inferior temporal sulcus. Although MSIT and code overlapped in frontal and parietal areas, like code with logic/math, the precise regions of overlap within these general locations differed.

Finally, code overlapped with language (language > math) in portions of the inferior frontal gyrus and the posterior aspect of the superior temporal sulcus/middle temporal gyrus. The overlap between language and code was on average low, and the degree of overlap varied considerably across participants (cosine sim range: [0.105, 0.480]), with only half of the participants showing above chance overlap. Notably there was no relationship between overlap of code and language and level of expertise, as measured either by years of experience coding (regression against code-language overlap: R^2^ = 0, p = 0.99; regression against code-math overlap: R^2^ = 0.033, p = 0.52) or performance on coding assessments (regression against code-language overlap: R^2^ = 0.033, p = 0.52; regression against code-math overlap: R^2^ = 0.064, p = 0.36).

#### Lateralization

The group activation map suggested that code comprehension is left-lateralized. Analyses of individual lateralization indices showed that indeed, code comprehension was as left-lateralized as language (Code lateralization index mean = 0.451, one-sample t-test against 0: t(14) = 5.501, p < 0.001; Language mean = 0.393, t(14) = 5.523, p < 0.001; paired t-test between code and language: t(14) = 1.203, p = 0.25). Moreover, lateralization indices of code and language were highly correlated across individuals (R^2^ = 0.658, p < 0.001) (Figure 5).

**Figure 5.**
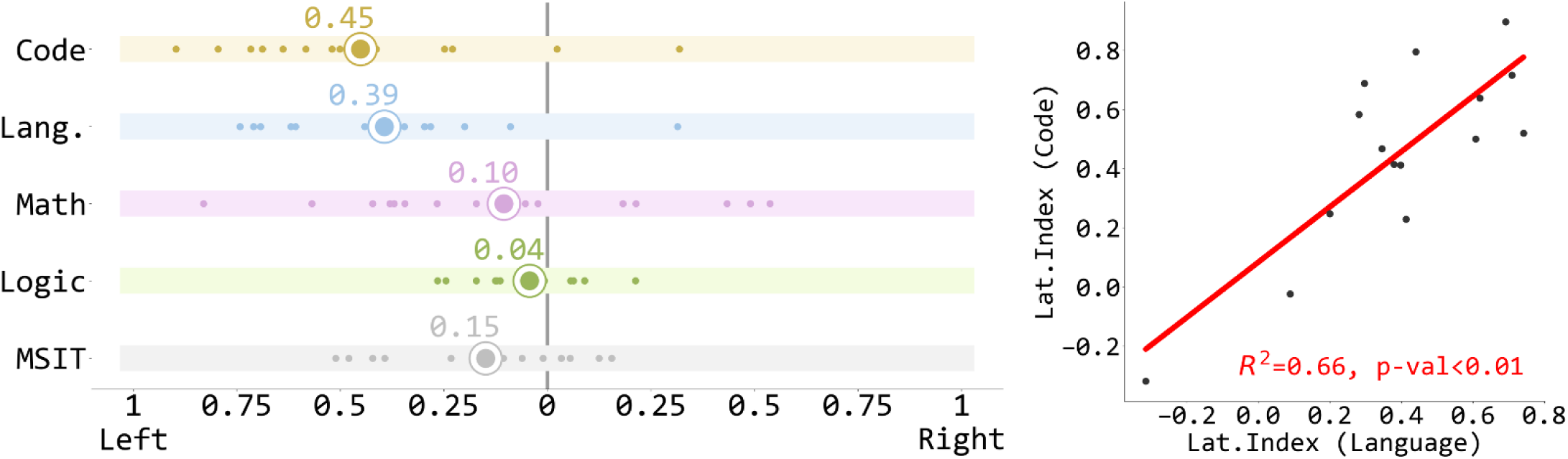
(a) The lateralization index of the code contrast and the localizer contrasts. Each white dot stands for one participant, and the enlarged dots represent the mean values. (b) The lateralization indices of code contrast and language contrast are highly correlated.

## Discussion

A consistent network of left-lateralized regions was activated across individuals during Python code comprehension. This network included the intraparietal sulcus (IPS), several regions within the lateral prefrontal cortex (PFC) and the posterior-inferior aspect of the middle temporal gyrus (pMTG). This code responsive network was more active during *real* than *fake code* trials, even though for expert Python coders, the *fake code* task was more difficult (as measured by reaction time) than the *real code* task. Within this code-responsive neural network, spatial patterns of activation distinguished between `for` vs. `if` code functions, suggesting that this network represents code-relevant information and is not merely activated during the coding task due to general difficulty demands. In overlap analyses, the code comprehension network was most similar to the fronto-parietal system involved in formal logical reasoning and to a lesser degree math. By contrast overlap with the perisylvian fronto-temporal language network is low.

### Code overlaps with logic

Code, logical reasoning, math and the MSIT task all activated aspects of the so-called fronto-parietal executive control system. However, overlap of code with logic was most extensive, followed by math and finally the MSIT. The difference between the MSIT task on the one hand and code comprehension, logic and math on the other, was particularly pronounced in the frontal lobe. There only code, logic and math activated more anterior regions of prefrontal cortex, including BA 46 and BA 9, although logic-associated activation extended even more anteriorly than code. These findings suggest that neural overlap between logic and code is specific, and not fully accounted for by the general involvement of the central executive system.

Previous studies also find that the fronto-parietal network, including anterior prefrontal areas, are involved in logical reasoning (Prado, Chadha, & Booth, 2011; Tsujii, Sakatani, Masuda, Akiyama, & Watanabe, 2011). For example, anterior PFC is active when participants solve formal logical problems with quantifiers (e.g. *<u>all</u> X are Y; Z is a X; therefore Z is Y*) and connectives (e.g. *<u>if</u> X then Y; not Y; therefore not X*) and plays a key role in deductive reasoning with variables (Coetzee & Monti, 2018; Goel, 2007; Goel & Dolan, 2004; Monti et al., 2009; Reverberi et al., 2010; Reverberi et al., 2007; Rodriguez-Moreno & Hirsch, 2009)

A fronto-parietal network has also been consistently implicated in math (Friedrich & Friederici, 2013; Maruyama, Pallier, Jobert, Sigman, & Dehaene, 2012; Piazza, Pinel, Le Bihan, & Dehaene, 2007; Wendelken, 2015). Some of the parietal responses to math have been linked to the processing of quantity information (Eger et al., 2009; Nieder, 2016; Nieder & Miller, 2004; Piazza & Eger, 2016; Roitman, Brannon, & Platt, 2007; Tudusciuc & Nieder, 2009). For example, neurons in the IPS of monkeys code numerosity of dots (Nieder, 2016). However, much of the same fronto-parietal network is also active during the processing of mathematical statements free of digits and arithmetic operations (Amalric & Dehaene, 2016, 2018; Wendelken, 2015). In the current study, both the anterior prefrontal areas and parietal areas involved in math also overlapped with code and logical reasoning. Some of this activation could therefore reflect common operations, such as the manipulation of rules and symbols in working memory. On the other hand, the lower overlap between coding and math as compared to overlap with coding and logic could be because only math involves quantitative processing.

The present evidence suggests that culturally derived symbol systems (i.e., code comprehension, formal logic and math) depend on a common fronto-parietal network, including the executive system. As noted in the introduction, although each of these symbol systems has its unique cognitive properties, they also have much in common. All involve the manipulation of abstract arbitrary symbols without inherent semantic content (e.g. `X, Y, input, result`) according to explicit rules. In the current logical inference and code experimental tasks, mental representations of several unknown variables are constructed (for logic “*X*”, “*Y*”, and “*Z*”, for code “`input`” and “`result`”) and the relationships between them deduced according to rules of formal logic or code.

There are also important differences between the rules of logical inference and programming. Take “if” conditional judgement for example again. In formal logic, the statement “*if P then Q*” doesn’t imply anything about what happens when *P* is false. On the contrary, in Python and most other programming languages, the statement

~~~
if condition==True:
      do_something()
~~~

automatically implies that when the condition is false, the function “`do_something()`” isn’t executed, unless otherwise specified. Learning to program involves acquiring the particular set of conventionalized rules used within programming languages and a syntax that specifies how the programming language in question expresses logical operations (Dalbey & Linn, 1985; Pea & Kurland, 1984; Pennington, 1987; Robins, Rountree, & Rountree, 2003). We speculate that such knowledge is encoded within the fronto-parietal network identified in the current study. Future studies comparing coders with different levels of expertise should test whether learning to code modifies circuits within the code-responsive neural network identified in the current study.

### The involvement of the multiple-demand executive control system in code comprehension

Code comprehension showed partial overlap with the MSIT task, particularly in the parietal cortex and in posterior frontal areas. Previous work has noted cognitive and neural similarity between arbitrary small-scale working memory tasks, such as the MSIT, and formal symbol systems (Anderson, 2005; Qin et al., 2004). As noted in the introduction, the MSIT task is a classic localizer for the executive function system (e.g., Stroop, n-back and MSIT) (Duncan, 2010; Fedorenko, Duncan, & Kanwisher, 2013; Miller & Cohen, 2001; Woolgar et al., 2011; Zanto & Gazzaley, 2013; Zhang et al., 2013). Like code comprehension, most experimental tasks that activate the central executive system involve the maintenance, manipulation and selection of arbitrary stimulus response mappings according to a set of predetermined rules (Woolgar et al., 2011; Zhang et al., 2013). For example, in the MSIT task among the many possible ways to map a visually presented digit triplet to a button press, the participants had to maintain in their working memory the rule to press the button whose index corresponds to the value of the unique digit in the triplet. In the difficult condition, participants use a less natural rule to make a response.

Previous studies also showed that the fronto-parietal executive system was involved in rule maintenance and switching, as well as variable representation. In one task-switching study the fronto-parietal executive system was active when participants maintained a cued rule in working memory and the level of activity increased with the complexity of the rule maintained (Bunge, Kahn, Wallis, Miller, & Wagner, 2003). Patterns of neural activity within the executive system encoded which rule is currently being applied and activity is modulated by rule switching (Buschman, Denovellis, Diogo, Bullock, & Miller, 2012; Crittenden & Duncan, 2014; Xu et al., 2017). Finally, studies with non-human primates found that neurons in the frontal lobe encode task-based variables (Duncan, 2010; Kennerley, Dahmubed, Lara, & Wallis, 2009; Nieder, 2013). Such processes, studied in the context of simple experimental tasks, may also play a role in code comprehension.

Although formal symbol systems and simple rule-based tasks share cognitive elements, tasks such as the MSIT involve simple rules that specify stimulus response mappings, rather than mental manipulations of variables. An intriguing possibility is that the neural machinery supporting code comprehension, as well as other culturally derived symbol systems, is a subset of a system that originally evolved for the maintenance and manipulation of simpler variables and rules (Anderson, 2005; Qin et al., 2004).

### Code comprehension and language

We find that the perisylvian fronto-temporal network that is selectively responsive to language, relative to math, does not overlap with the neural network involved in code comprehension. Previous studies also found that math and formal logic did not depend on classic language networks (Amalric & Dehaene, 2016; Monti et al., 2009). Lack of overlap between code and language is intriguing given the cognitive similarities between these domains (Fedorenko et al., 2019; Pandža, 2016; Peitek et al., 2018; Portnoff, 2018; Prat et al., 2020; Siegmund et al., 2014). As noted in the introduction, programming languages borrow letters and words from natural language, and both natural language and code have hierarchical, recursive grammars (Fitch et al., 2005).

One possible explanation for low overlap between the perisylvian fronto-temporal language network and code is that the language system is evolutionarily predisposed to support natural language processing and not generalizable even to similar domains, like computer code and formal logic (Dehaene-Lambertz, Hertz-Pannier, & Dubois, 2006; Fedorenko et al., 2011). Timing could also play a role. The perisylvian fronto-temporal language network may have a sensitive period of development during which it is most capable of learning (Cheng, Roth, Halgren, & Mayberry, 2019; Mayberry, Davenport, Roth, & Halgren, 2018; Ramirez et al., 2016). By the time people learn to code, the network may be incapable of taking on new cognitive functions. Indeed, even acquiring a second language late in life leads to lower levels of proficiency and responses outside the perisylvian fronto-temporal system (Hartshorne, Tenenbaum, & Pinker, 2018; Johnson & Newport, 1989). These observations suggest that domain-specific systems, like the perisylvian fronto-temporal language network, are not always amenable for “recycling” by cultural inventions. The fronto-parietal system might be inherently more flexible throughout the lifespan and thus more capable of taking on new cultural skills (Riley, Qi, Zhou, & Constantinidis, 2018).

Despite lack of direct overlap, lateralization patterns of language and coding were highly correlated across individuals i.e. those individuals with highly left-lateralized responses to sentences also showed highly left lateralized responses to code. This intriguing observation suggests that the relationship between code and language may be ontogenetic as well as phylogenetic. It is hard to imagine how code in its current form could have been invented in the absence of language (Fitch et al., 2005). Ontogenetically, code-relevant neural representations might be enabled by the language system, even though they are distinct from it.

An analogous example comes from the domain of reading (Dehaene et al., 2010; McCandliss, Cohen, & Dehaene, 2003). Reading-relevant regions, such as the visual word form area (VWFA), are strongly co-lateralized with the perisylvian fronto-temporal language network across people (Cai, Paulignan, Brysbaert, Ibarrola, & Nazir, 2010). The VWFA has strong anatomical connectivity with the fronto-temporal language network even prior to literacy (Bouhali et al., 2014; Saygin et al., 2016). Analogously, code comprehension may colonize a left-lateralized portion of the central executive system due to its stronger (i.e., within hemisphere) connectivity with the perisylvian fronto-temporal language network.

## Conclusions

A fronto-parietal cortical network is consistently engaged in expert programmers during code comprehension. Patterns of activity within this network distinguish between `for` and `if` functions. This network overlaps with other culturally derived symbol systems, in particular formal logic and to a lesser degree math. By contrast, the neural basis of code is distinct from the perisylvian fronto-temporal language network. Rather than recycling domain specific cortical mechanisms for language, code, like formal logic and math, depends on a subset of the domain general executive system, including anterior prefrontal areas. The executive system may be uniquely suited as a flexible learning mechanism capable of supporting an array of cultural symbol systems acquired in adulthood.

## Supporting information

supplementary methods and results

## Notes

### Competing Interest Statement

The authors have declared no competing interest.

### Summary of Updates

separate supplementary material as individual files

