## supplementary methods and results for "Computer code comprehension shares neural resources with formal logical inference in the fronto-parietal network"

An example of the difficult out-of-the-scanner exercise

```
Consider the following code:
###

def xx(a):
    b = a+3
    return
if xx(1)==________:
    print("HEY!!")

###
What keyword should be filled in the blank if we want the code to print "HEY!!"?
(It's a keyword. There should be only letters in your answer.)
```

Detailed information about the stimuli

Across all real functions, the variable names input, result, and ii, were the same, and only the 12 built-in functions (capitalize(), isalnum(), isalpha(), isdigit(), len(), lower(), range(), sorted(), split(), str(), swapcase(), and upper()) and 3 expressions (list comprehension, slice notation, and string formatting, see Table S1 for examples) tested during the screening exercise were included in the user-defined functions.

The addition operator (+) occurred in all functions, but always meant string concatenation rather than numeric addition, and never took a numeric as operand. In each group, the multiplication operator (*) existed in 32 out of the 96 real functions, and 10 of them took a numeric as one of its operands. However, in all these instances, the “multiplication” meant repetition of strings or lists instead of numeric multiplications (e.g. “abc”*3 results in “abcabcabc”). In each group, 12 out of the 96 real functions contained a comparison to a numeric value, such as “len(input) > 5”.

The two variants of each control structure

We designed two variants to implement each control structure, for and if. In the first variant of a for code, the for loop was implemented in the canonical way. In the second variant of a for code, we implemented the loop with a Python-specific expression “list comprehension”, where the operation to be performed on each element in a list (or string) was stated before specifying the list to be iterated over. In the first variant of an if code, the if conditional was implemented in the canonical way. In the second variant of an if code, the conditional was implemented by first stating the action to take if a condition is true, then multiplying this action to the true/false judgement statement of the condition. There was not a formal jargon for this kind of implementation, for the sake of convenience, we called it “conditional multiplication” in this study. Please refer to Table S1 for examples of each variant.

The algorithm of the permutation of stimuli

In this experiment, there were 5 conditions, “FOR1”, “FOR2”, “IF1”, “IF2”, and “FAKE”. For simplicity, from here on we label them as “A”, “B”, “C”, “D”, and “E”, respectively.

There were 120 permutations for 5 distinct labels, such as “ABCDE”, “BCDEA”, “CDEAB”, “DEABC”, “EABCD”, “ACBDE”, “CBDEA”, etc. Each run consisted of 20 functions, which was 4 permutations of 5 labels. Therefore, for each run, we drew 4 permutations out of the 120 possible permutations. So, the order a participant saw in the first run can be:

*ABCDE BCDEA CDEAB DEABC*

And the order in the second run can be:

*EABCD ACBDE CBDEA BDEAC*

The permutations were allocated such that every participant saw 24 permutations across all 6 runs, and every 5 participants saw all the 120 permutations.

After determining the order of the conditions, we assigned actual instances of the conditions to the labels. The order of presentation for the functions in each condition was also permuted such that a function in run 1 for participant 1 appeared in run 2 for participant 2, run 6 for participant 6, run 1 for participant 7, and so on. Specifically, the first run of the first participant could be:

*A_1_B_1_C_1_D_1_E_1_ B_2_C_2_D_2_E_2_A_2_ C_3_D_3_E_3_A_3_B_3_ D_4_E_4_A_4_B_4_C_4_*

While the first run of the second participant could be:

*A_5_B_5_C_5_D_5_E_5_ B_6_C_6_D_6_E_6_A_6_ C_7_D_7_E_7_A_7_B_7_ D_8_E_8_A_8_B_8_C_8_*

The second participant still saw *A_1_, B_1_, C_1_, …… D_4,_ E_4_*, just in some later runs.

As a result of permutations of both conditions and functions within condition, all of the participants saw a unique order of presentation.

### Supplementary results

Overlap analysis in the right hemisphere

Code comprehension (real > fake) overlapped significantly above chance with all localizer tasks: logic, math, language and MSIT (each task compared to chance p’s < 10-6 compared to code split-half overlap p’s < 10-6). The degree of overlap differed significantly across tasks (repeated-measures ANOVA: F(3,42) = 3.03, p = 0.040). Post-hoc paired t-tests (FDR-corrected p’s < 0.05) revealed that the overlap with logic was significantly higher than with math and MSIT, but indistinguishable from the overlap with language. The overlaps with the other there tasks (language, math, MSIT) were statistically indistinguishable from each other (Table S3).
