## supplementary table S1 for "Computer code comprehension shares neural resources with formal logical inference in the fronto-parietal network"

Table S1. All the functions included in the fMRI study

| **Real / Fake** | **Control structure** | | **Variant** | **Code** |
| --- | --- | --- | --- | --- |
| real | | FOR | Canonical | def fun(input):  result = []  for ii in input.split("."):  result += [ii.isalpha()]  return result |
| real | | FOR | Canonical | def fun(input):  result = input.split(".")  for ii in range(len(result)):  result[ii] += "..."  return result |
| real | | FOR | Canonical | def fun(input):  result = []  for ii in input.split("."):  result += [ii.capitalize()]  return result |
| real | | FOR | Canonical | def fun(input):  result=[]  for ii in range(len(input)):  result+=["%d:%s"%(ii,input[ii])]  return result |
| real | | FOR | Canonical | def fun(input):  result = input.split("_")  for ii in range(len(result)):  result[ii] = result[ii]*2  return result |
| real | | FOR | Canonical | def fun(input):  result=["input"]  for ii in input:  result += ["=%s"%ii]  return result |
| real | | FOR | Canonical | def fun(input):  result = []  for ii in input.split("."):  result += [ii.isalpha()]  return result |
| real | | FOR | Canonical | def fun(input):  result = input.split("-")  for ii in range(len(result)):  result[ii] += ":"  return result |
| real | | FOR | Canonical | def fun(input):  result = ["result:"]  for ii in input:  result += ["%s's"%ii]  return result |
| real | | FOR | Canonical | def fun(input):  result = input.split(" ")  for ii in range(len(result)):  result[ii] = result[ii][0]  return result |
| real | | FOR | Canonical | def fun(input):  result=[]  for ii in input:  result += ["%s+%s"%(ii,ii)]  return result |
| real | | FOR | Canonical | def fun(input):  result = ["for"]  for ii in input:  result += [ii]  return result |
| real | | FOR | Canonical | def fun(input):  result = ["ii+ii"]  for ii in input:  result += [ii*2]  return result |
| real | | FOR | Canonical | def fun(input):  result = ["for ii:"]  for ii in input:  result += ["/"+ii+"/"]  return result |
| real | | FOR | Canonical | def fun(input):  result=[input]  for ii in range(len(input)):  result+=["=" + "_" + "="]  return result |
| real | | FOR | Canonical | def fun(input):  result=["%"]  for ii in input:  result += ["%s,"%ii]  return result |
| real | | FOR | Canonical | def fun(input):  result=["%%"]  for ii in range(len(input)):  result+=[str(ii)+"%"]  return result |
| real | | FOR | Canonical | def fun(input):  result = [input]  for ii in input.split("."):  result += [ii[0].upper()]  return result |
| real | | FOR | Canonical | def fun(input):  result=["..."]  for ii in range(len(input)):  result+=[input[ii]]  return result |
| real | | FOR | Canonical | def fun(input):  result = ["in:"]  for ii in input:  result += [ii+":"]  return result |
| real | | FOR | Canonical | def fun(input):  result = input.split("+")  for ii in range(len(result)):  result[ii] += "-"  return result |
| real | | FOR | Canonical | def fun(input):  result = ["="]  for ii in input.split("="):  result += [ii.swapcase()]  return result |
| real | | FOR | Canonical | def fun(input):  result=["in"]  for ii in range(len(input)):  result+=["%d%d"%(ii,ii)]  return result |
| real | | FOR | Canonical | def fun(input):  result = ["input"]  for ii in input.split("_"):  result += [ii[0].lower()]  return result |
| real | | FOR | Canonical | def fun(input):  result = ["result="]  for ii in input:  result += [ii*2]  return result |
| real | | FOR | Canonical | def fun(input):  result = []  for ii in input.split("-"):  result += [ii.upper()]  return result |
| real | | FOR | Canonical | def fun(input):  result=[]  for ii in range(len(input)):  result+=["%d-->%.3d"%(ii,ii)]  return result |
| real | | FOR | Canonical | def fun(input):  result=[".."]  for ii in input:  result += ["%s!!"%ii]  return result |
| real | | FOR | Canonical | def fun(input):  result=[]  for ii in range(len(input)):  result+=["[%s]"%input[ii]]  return result |
| real | | FOR | Canonical | def fun(input):  result = input.split("/")  for ii in range(len(result)):  result[ii] += "->"  return result |
| real | | FOR | Canonical | def fun(input):  result = ["for:"]  for ii in input.split(","):  result += [ii.lower()]  return result |
| real | | FOR | Canonical | def fun(input):  result=["for"]  for ii in input:  result += ["in %s"%ii]  return result |
| real | | FOR | Canonical | def fun(input):  result = input.split("*")  for ii in range(len(result)):  result[ii] += "a"  return result |
| real | | FOR | Canonical | def fun(input):  result = [input]  for ii in input:  result += [ii+ii]  return result |
| real | | FOR | Canonical | def fun(input):  result=[input]  for ii in input:  result += ["%s"%ii]  return result |
| real | | FOR | Canonical | def fun(input):  result=[]  for ii in range(len(input)):  result+=["%s++"%str(ii)]  return result |
| real | | FOR | Canonical | def fun(input):  result = ["input"]  for ii in input:  result += [ii+"."]  return result |
| real | | FOR | Canonical | def fun(input):  result = []  for ii in input.split("_"):  result += [ii.lower()]  return result |
| real | | FOR | Canonical | def fun(input):  result=[input]  for ii in range(len(input)):  result+=["%d"%ii]  return result |
| real | | FOR | Canonical | def fun(input):  result = input.split("*")  for ii in range(len(result)):  result[ii] = None  return result |
| real | | FOR | Canonical | def fun(input):  result = []  for ii in input:  result += ["("+ii+")"]  return result |
| real | | FOR | Canonical | def fun(input):  result = []  for ii in input.split("%"):  result += [ii.isdigit()]  return result |
| real | | FOR | Canonical | def fun(input):  result=[input]  for ii in input:  result += ["[%s]"%ii]  return result |
| real | | FOR | Canonical | def fun(input):  result = input.split(".")  for ii in range(len(result)):  result[ii] = result[ii][-2:]  return result |
| real | | FOR | Canonical | def fun(input):  result = [input]  for ii in input.split("="):  result += [ii[0].lower()]  return result |
| real | | FOR | Canonical | def fun(input):  result = input.split(".")  for ii in range(len(result)):  result[ii] += "."  return result |
| real | | FOR | Canonical | def fun(input):  result=[]  for ii in range(len(input)):  result+=["[%s]"%input[-1]]  return result |
| real | | FOR | Canonical | def fun(input):  result=[]  for ii in input:  result += ["%s"%(ii+"s")]  return result |
| real | | FOR | List comprehension | def fun(input):  result = [".."]  result += ["%s!!"%ii  for ii in input]  return result |
| real | | FOR | List comprehension | def fun(input):  result = [ "for", "ii" ]  result += [ii + ii  for ii in input]  return result |
| real | | FOR | List comprehension | def fun(input):  result = input.split(".")  result = [ii + "://" for  ii in result]  return result |
| real | | FOR | List comprehension | def fun(input):  result=[]  result+=["%s++"%str(ii)  for ii in range(len(input))]  return result |
| real | | FOR | List comprehension | def fun(input):  result = ["input"]  result += [ii + "."  for ii in input]  return result |
| real | | FOR | List comprehension | def fun(input):  result = []  result += [ii.lower()  for ii in input.split("_")]  return result |
| real | | FOR | List comprehension | def fun(input):  result=[]  result+=["%d-->%.3d"%(ii,ii)  for ii in range(len(input))]  return result |
| real | | FOR | List comprehension | def fun(input):  result=[input]  result+=["%d"%ii  for ii in range(len(input))]  return result |
| real | | FOR | List comprehension | def fun(input):  result = input.split("*")  result = [None for  ii in result]  return result |
| real | | FOR | List comprehension | def fun(input):  result=[]  result+=["[%s]"%input[ii]  for ii in range(len(input))]  return result |
| real | | FOR | List comprehension | def fun(input):  result = [ "result=" ]  result += [ii*2  for ii in input]  return result |
| real | | FOR | List comprehension | def fun(input):  result = []  result += [ii.upper()  for ii in input.split("-")]  return result |
| real | | FOR | List comprehension | def fun(input):  result = [input]  result += ["[%s]"%ii  for ii in input]  return result |
| real | | FOR | List comprehension | def fun(input):  result = []  result = [ii.isdigit()  for ii in input.split("%")]  return result |
| real | | FOR | List comprehension | def fun(input):  result = input.split("/")  result = [result[ii]+"->" for  ii in range(len(result))]  return result |
| real | | FOR | List comprehension | def fun(input):  result = [ "for:" ]  result += [ii.lower()  for ii in input.split(",")]  return result |
| real | | FOR | List comprehension | def fun(input):  result = input.split(".")  result = [ii[-2:] for  ii in result]  return result |
| real | | FOR | List comprehension | def fun(input):  result = []  result += ["(" + ii + ")"  for ii in input]  return result |
| real | | FOR | List comprehension | def fun(input):  result = input.split("*")  result = [result[ii]+"a" for  ii in range(len(result))]  return result |
| real | | FOR | List comprehension | def fun(input):  result = [ "for" ]  result += ["in %s"%ii  for ii in input]  return result |
| real | | FOR | List comprehension | def fun(input):  result = []  result += ["%s"%(ii + "s")  for ii in input]  return result |
| real | | FOR | List comprehension | def fun(input):  result = [input]  result += ["%s"%ii  for ii in input]  return result |
| real | | FOR | List comprehension | def fun(input):  result=[]  result+=["[%s]"%(input[-1])  for ii in range(len(input))]  return result |
| real | | FOR | List comprehension | def fun(input):  result = [input]  result += [ii + ii  for ii in input]  return result |
| real | | FOR | List comprehension | def fun(input):  result=[]  result+=["%d:%s"%(ii,input[ii])  for ii in range(len(input))]  return result |
| real | | FOR | List comprehension | def fun(input):  result = []  result = [ii.capitalize()  for ii in input.split(".")]  return result |
| real | | FOR | List comprehension | def fun(input):  result = []  result = [ii.isalpha()  for ii in input.split(".")]  return result |
| real | | FOR | List comprehension | def fun(input):  result = input.split(".")  result = [ii+"..." for  ii in result]  return result |
| real | | FOR | List comprehension | def fun(input):  result = []  result = [ii.isalpha()  for ii in input.split(".")]  return result |
| real | | FOR | List comprehension | def fun(input):  result = input.split("_")  result = [ii*2 for  ii in result]  return result |
| real | | FOR | List comprehension | def fun(input):  result = input.split("-")  result = [ii + ":" for  ii in result]  return result |
| real | | FOR | List comprehension | def fun(input):  result = ["input"]  result += ["=%s"%ii  for ii in input]  return result |
| real | | FOR | List comprehension | def fun(input):  result = ["for"]  result += [ii  for ii in input]  return result |
| real | | FOR | List comprehension | def fun(input):  result = []  result += ["%s+%s"%(ii, ii)  for ii in input]  return result |
| real | | FOR | List comprehension | def fun(input):  result = [ "result:" ]  result += ["%s's"%ii  for ii in input]  return result |
| real | | FOR | List comprehension | def fun(input):  result = input.split(" ")  result = [result[ii][0] for  ii in range(len(result))]  return result |
| real | | FOR | List comprehension | def fun(input):  result=[input]  result+=["=" + "_" + "="  for ii in range(len(input))]  return result |
| real | | FOR | List comprehension | def fun(input):  result = [ "for ii:" ]  result += ["/" + ii + "/"  for ii in input]  return result |
| real | | FOR | List comprehension | def fun(input):  result = [ "%" ]  result += ["%s,"%ii  for ii in input]  return result |
| real | | FOR | List comprehension | def fun(input):  result = ["ii+ii"]  result += [ii*2  for ii in input]  return result |
| real | | FOR | List comprehension | def fun(input):  result=["..."]  result+=[input[ii] for ii  in range(len(input))]  return result |
| real | | FOR | List comprehension | def fun(input):  result = ["in:"]  result += [ii + ":"  for ii in input]  return result |
| real | | FOR | List comprehension | def fun(input):  result = [ input ]  result += [ii[0].upper()  for ii in input.split(".")]  return result |
| real | | FOR | List comprehension | def fun(input):  result=["%%"]  result+=[str(ii) + "%"  for ii in range(len(input))]  return result |
| real | | FOR | List comprehension | def fun(input):  result = input.split("+")  result = [ii + "-" for  ii in result]  return result |
| real | | FOR | List comprehension | def fun(input):  result = ["input"]  result += [ii[0].lower()  for ii in input.split("_")]  return result |
| real | | FOR | List comprehension | def fun(input):  result=["in"]  result+=["%d%d"%(ii, ii) for  ii in range(len(input))]  return result |
| real | | FOR | List comprehension | def fun(input):  result = ["="]  result += [ii.swapcase()  for ii in input.split("=")]  return result |
| real | | IF | Canonical | def fun(input):  result = "input = "  if input[0]!=".":  result += input.upper()  return result |
| real | | IF | Canonical | def fun(input):  result = "not digit "  if not input.isdigit():  result += result*2  return result |
| real | | IF | Canonical | def fun(input):  result = "..."  if len(input.split("."))>1:  result += input.split(".")[-1]  return result |
| real | | IF | Canonical | def fun(input):  result = "is digit."  if input.isdigit():  result += result*2  return result |
| real | | IF | Canonical | def fun(input):  result=["result:"]  if not input.isalpha():  result+=sorted(input)  return result |
| real | | IF | Canonical | def fun(input):  result = "result: "  if input[-1]!=".":  result += input[-1].upper()  return result |
| real | | IF | Canonical | def fun(input):  result = "result: "  if input[-1]=="j":  result += input[-1].upper()  return result |
| real | | IF | Canonical | def fun(input):  result = input[:-5]  if input.isdigit():  result += result*4  return result |
| real | | IF | Canonical | def fun(input):  result = input[0]  if not input.isalpha():  result += result.upper()  return result |
| real | | IF | Canonical | def fun(input):  result=["input:"]  if len(input)<5:  result+=sorted(input)  return result |
| real | | IF | Canonical | def fun(input):  result=[]  if not input.isalnum():  result+=sorted(input)  return result |
| real | | IF | Canonical | def fun(input):  result = "result: "  if input[-1]==".":  result += input[-1].upper()  return result |
| real | | IF | Canonical | def fun(input):  result = [input]  if not input.isalpha():  result += [input]  return result |
| real | | IF | Canonical | def fun(input):  result = "split: "  if len(input.split("+"))<3:  result += input  return result |
| real | | IF | Canonical | def fun(input):  result=[":"]  if input.isdigit():  result+=sorted(input)  return result |
| real | | IF | Canonical | def fun(input):  result = "input"  if input.isalpha():  result += input.upper()  return result |
| real | | IF | Canonical | def fun(input):  result = ["input"]  if len(input.split("."))<4:  result += input.split(".")  return result |
| real | | IF | Canonical | def fun(input):  result = input[:-1]  if not input[-1].isdigit():  result += input[-1]  return result |
| real | | IF | Canonical | def fun(input):  result = [input]  if input.isdigit():  result += [len(input)]  return result |
| real | | IF | Canonical | def fun(input):  result = input  if len(input.split("/"))>1:  result += input.split("/")[-1]  return result |
| real | | IF | Canonical | def fun(input):  result = "digit..."  if not input.isdigit():  result += input*2  return result |
| real | | IF | Canonical | def fun(input):  result = input[:-1]  if input[-1]!="a":  result += input[-1].upper()  return result |
| real | | IF | Canonical | def fun(input):  result = "split+"  if len(input.split(">"))>1:  result += input.split(">")[0]  return result |
| real | | IF | Canonical | def fun(input):  result = "if is "  if input.isalpha():  result += "alpha"  return result |
| real | | IF | Canonical | def fun(input):  result = "result: "  if input[0]=="a":  result += input[0].upper()  return result |
| real | | IF | Canonical | def fun(input):  result=["result:"]  if input.isalnum():  result+=sorted(input)  return result |
| real | | IF | Canonical | def fun(input):  result = ">>> "  if len(input.split("."))>1:  result += input.split(".")[0]  return result |
| real | | IF | Canonical | def fun(input):  result = "i"  if input.isalpha():  result += input.upper()  return result |
| real | | IF | Canonical | def fun(input):  result = "input: "  if not input.isdigit():  result += input  return result |
| real | | IF | Canonical | def fun(input):  result = "..."  if len(input.split("-"))>1:  result += input.split("-")[-1]  return result |
| real | | IF | Canonical | def fun(input):  result = "i"  if input.isalpha():  result += input.lower()  return result |
| real | | IF | Canonical | def fun(input):  result = "."  if not input.isalpha():  result += input  return result |
| real | | IF | Canonical | def fun(input):  result = input  if len(input.split("="))>1:  result += input.split("=")[0]  return result |
| real | | IF | Canonical | def fun(input):  result = input  if not input.isdigit():  result += input[-1]  return result |
| real | | IF | Canonical | def fun(input):  result = "input"  if input.isdigit():  result += result*3  return result |
| real | | IF | Canonical | def fun(input):  result = "[input] "  if input[0]!="_":  result += input.capitalize()  return result |
| real | | IF | Canonical | def fun(input):  result = "result: "  if len(input.split("."))>1:  result += input.split(".")[0]  return result |
| real | | IF | Canonical | def fun(input):  result="result:"  if not input.isdigit():  result+=sorted(input)[0]  return result |
| real | | IF | Canonical | def fun(input):  result = "result: "  if input[0]=="r":  result += input.capitalize()  return result |
| real | | IF | Canonical | def fun(input):  result = "(isalpha)"  if input.isalpha():  result += result*2  return result |
| real | | IF | Canonical | def fun(input):  result = "[result] "  if input[0]!="_":  result += input[0]*2  return result |
| real | | IF | Canonical | def fun(input):  result=[input[:-1]]  if input.isalpha():  result+=sorted(input[:-1])  return result |
| real | | IF | Canonical | def fun(input):  result = "h"  if not input.isalpha():  result += input.upper()  return result |
| real | | IF | Canonical | def fun(input):  result=[input]  if len(input)<=3:  result+=sorted(input)  return result |
| real | | IF | Canonical | def fun(input):  result = "result: "  if input.isdigit():  result += input.lower()  return result |
| real | | IF | Canonical | def fun(input):  result = ["result"]  if len(input.split("_"))<3:  result += input.split("_")  return result |
| real | | IF | Canonical | def fun(input):  result = "input: "  if input[0]==".":  result += input.swapcase()  return result |
| real | | IF | Canonical | def fun(input):  result=["sorted"]  if input.isalpha():  result+=sorted(input)  return result |
| real | | IF | Conditional multiplication | def fun(input):  result = "result: "  result += (input[0]  .upper()*(input[0]=="a"))  return result |
| real | | IF | Conditional multiplication | def fun(input):  result=["result:"]  result+=(input  .isalnum()*sorted(input))  return result |
| real | | IF | Conditional multiplication | def fun(input):  result = ">>> "  result += (input.split(".")[0]  *(len(input.split("."))>1))  return result |
| real | | IF | Conditional multiplication | def fun(input):  result = "i"  result += (input  .upper()*input.isalpha())  return result |
| real | | IF | Conditional multiplication | def fun(input):  result = "input: "  result += (input*  (not input.isdigit()))  return result |
| real | | IF | Conditional multiplication | def fun(input):  result = "..."  result += (input.split("-")[-1]  *(len(input.split("-"))>1))  return result |
| real | | IF | Conditional multiplication | def fun(input):  result = "."  result += (input*  (not input.isalpha()))  return result |
| real | | IF | Conditional multiplication | def fun(input):  result = "i"  result += (input  .lower()*input.isalpha())  return result |
| real | | IF | Conditional multiplication | def fun(input):  result = input  result += (input[-1]*  (not input.isdigit()))  return result |
| real | | IF | Conditional multiplication | def fun(input):  result = "[input] "  result += (input  .capitalize()*(input[0]!="_"))  return result |
| real | | IF | Conditional multiplication | def fun(input):  result = input  result += (input.split("=")[0]  *(len(input.split("="))>1))  return result |
| real | | IF | Conditional multiplication | def fun(input):  result = "input"  result += (result*  3*input.isdigit())  return result |
| real | | IF | Conditional multiplication | def fun(input):  result = "result: "  result += (input.split(".")[0]  *(len(input.split("."))>1))  return result |
| real | | IF | Conditional multiplication | def fun(input):  result = "(isalpha)"  result += (result*  2*input.isalpha())  return result |
| real | | IF | Conditional multiplication | def fun(input):  result = "result: "  result += (input  .capitalize()*(input[0]=="r"))  return result |
| real | | IF | Conditional multiplication | def fun(input):  result="result:"  result+=(sorted(input)[0]*  (not input.isdigit()))  return result |
| real | | IF | Conditional multiplication | def fun(input):  result = "[result] "  result += (input[0]*2  *(input[0]!="_"))  return result |
| real | | IF | Conditional multiplication | def fun(input):  result=[input]  result+=(sorted(input)*  (len(input)<=3))  return result |
| real | | IF | Conditional multiplication | def fun(input):  result = "h"  result += (input.upper()*  (not input.isalpha()))  return result |
| real | | IF | Conditional multiplication | def fun(input):  result=[input[:-1]]  result+=(input  .isalpha()*sorted(input[:-1]))  return result |
| real | | IF | Conditional multiplication | def fun(input):  result = ["result"]  result += (input.split("_")  *(len(input.split("_"))<3))  return result |
| real | | IF | Conditional multiplication | def fun(input):  result = "result: "  result += (input.lower()  *input.isdigit())  return result |
| real | | IF | Conditional multiplication | def fun(input):  result=["sorted"]  result+=(input  .isalpha()*sorted(input))  return result |
| real | | IF | Conditional multiplication | def fun(input):  result = "input: "  result += (input  .swapcase()*(input[0]=="."))  return result |
| real | | IF | Conditional multiplication | def fun(input):  result = "..."  result += (input.split(".")[-1]  *(len(input.split("."))>1))  return result |
| real | | IF | Conditional multiplication | def fun(input):  result = "is digit."  result += (result*  2*input.isdigit())  return result |
| real | | IF | Conditional multiplication | def fun(input):  result = "input = "  result += (input  .upper()*(input[0]!="."))  return result |
| real | | IF | Conditional multiplication | def fun(input):  result = "not digit "  result += (result*  2*(not input.isdigit()))  return result |
| real | | IF | Conditional multiplication | def fun(input):  result=["result:"]  result+=(sorted(input)*  (not input.isalpha()))  return result |
| real | | IF | Conditional multiplication | def fun(input):  result = "result: "  result += (input[-1]  .upper()*(input[-1]!="."))  return result |
| real | | IF | Conditional multiplication | def fun(input):  result = input[:-5]  result += (result*  4*input.isdigit())  return result |
| real | | IF | Conditional multiplication | def fun(input):  result = "result: "  result += (input[-1]  .upper()*(input[-1]=="j"))  return result |
| real | | IF | Conditional multiplication | def fun(input):  result=["input:"]  result+=(sorted(input)*  (len(input)<5))  return result |
| real | | IF | Conditional multiplication | def fun(input):  result = input[0]  if not input.isalpha():  result += result.upper()  return result |
| real | | IF | Conditional multiplication | def fun(input):  result = "result: "  result += (input[-1]  .upper()*(input[-1]=="."))  return result |
| real | | IF | Conditional multiplication | def fun(input):  result=[]  result+=((not input  .isalnum())*sorted(input))  return result |
| real | | IF | Conditional multiplication | def fun(input):  result = "split: "  result += (input*  (len(input.split("+"))<3))  return result |
| real | | IF | Conditional multiplication | def fun(input):  result = [input]  result += ([input]*  (not input.isalpha()))  return result |
| real | | IF | Conditional multiplication | def fun(input):  result=[":"]  result+=(input  .isdigit()*sorted(input))  return result |
| real | | IF | Conditional multiplication | def fun(input):  result = "input"  result += (input  .upper()*input.isalpha())  return result |
| real | | IF | Conditional multiplication | def fun(input):  result = ["input"]  result += (input.split(".")  *(len(input.split("."))<4))  return result |
| real | | IF | Conditional multiplication | def fun(input):  result = input[:-1]  result += (input[-1]*  (not input[-1].isdigit()))  return result |
| real | | IF | Conditional multiplication | def fun(input):  result = input  result += (input.split("/")[-1]  *(len(input.split("/"))>1))  return result |
| real | | IF | Conditional multiplication | def fun(input):  result = [input]  result += ([len(input)]  *input.isdigit())  return result |
| real | | IF | Conditional multiplication | def fun(input):  result = input[:-1]  result += (input[-1]  .upper()*(input[-1]!="a"))  return result |
| real | | IF | Conditional multiplication | def fun(input):  result = "if is "  result += ("alpha"*  input.isalpha())  return result |
| real | | IF | Conditional multiplication | def fun(input):  result = "digit..."  result += (input*  2*(not input.isdigit()))  return result |
| real | | IF | Conditional multiplication | def fun(input):  result = "split+"  result += (input.split(">")[0]  *(len(input.split(">"))>1))  return result |
| fake | | N/A | N/A | input :def)fun(  result [= ]  input" in) split.for" ii(:.  isalpha[ii] result+.( )=  result return |
| fake | | N/A | N/A | input :def)fun(  result [= ]  input" in) split.for" ii(:.  isalpha[ii] result+.( )=  result return |
| fake | | N/A | N/A | input :def)fun(  split(input) result=". ".  range) for: in(result)len (ii  result. ii" +.".]=[  result return |
| fake | | N/A | N/A | input :def)fun(  result[=]  range) for: in(input)len (ii  ii]result:s"ii,d%input=[]")[+%%(  result return |
| fake | | N/A | N/A | input :def)fun(  result=input""[]  for: input ii in  result]s%ii+=" [= %"  result return |
| fake | | N/A | N/A | input :def)fun(  result [= ]  input" in) split.for" ii(:.  capitalize[ii] result+.( )=  result return |
| fake | | N/A | N/A | input :def)fun(  split(input) result="- ".  range) for: in(result)len (ii  result:ii]+= [" "  result return |
| fake | | N/A | N/A | input :def)fun(  split(input) result=". ".  range) for: in(result)len (ii  result/ ii" +/":]=[  result return |
| fake | | N/A | N/A | input :def)fun(  result"ii" for,"]"= [  ii for in: input  ii] result [ii=++  result return |
| fake | | N/A | N/A | input :def)fun(  result[=]  range) for: in(input)len (ii  s"ii[result+str%+)%"=+](  result return |
| fake | | N/A | N/A | input :def)fun(  result"=]"[..  ii for in: input  s" result!ii!=%"%] +[  result return |
| fake | | N/A | N/A | input :def)fun(  result=input" "[ ]  for: input ii in  result[ii+ "]+" .=  result return |
| fake | | N/A | N/A | input :def)fun(  result[=]  range) for: in(input)len (ii  ii]result-3d,ii-d%"=>[")(+%%.  result return |
| fake | | N/A | N/A | input :def)fun(  result=input" "[ ]  result .ii= [+ +" "  ]input for in ii  result return |
| fake | | N/A | N/A | input :def)fun(  result " ii" for,"] "= [  result [ ii =ii+ +  for in ]ii input  result return |
| fake | | N/A | N/A | input :def)fun(  result" = ]"[..  result%s"ii+%" [= !!  for in ]ii input  result return |
| fake | | N/A | N/A | input :def)fun(  result[=]  result[str+s=ii)"%+"(+%  range) for] in(input)len (ii  result return |
| fake | | N/A | N/A | input :def)fun(  split(input) result=". ".  result+ for / ii:/ ""= [  in ii ]result  result return |
| fake | | N/A | N/A | input :def)fun(  result[=]  3d)d%result%ii=ii-+[>-%.,(""  range) for] in(input)len (ii  result return |
| fake | | N/A | N/A | input :def)fun(  result=input" "[ ]  result[ii" s=%%"+ =  ]input for in ii  result return |
| fake | | N/A | N/A | input :def)fun(  result [= ]  capitalize) result .ii[=(  input" in) split.for" ii(].  result return |
| fake | | N/A | N/A | input :def)fun(  split(input) result=". ".  result+for. ii.. ""= [  in ii] result  result return |
| fake | | N/A | N/A | input :def)fun(  result [= ]  isalpha) result .ii[=(  input" in) split.for" ii(].  result return |
| fake | | N/A | N/A | input :def)fun(  result[=]  s)input[d%result=ii:ii+["%%(],"  range) for] in(input)len (ii  result return |
| fake | | N/A | N/A | input :def)fun(  split(input) result="- ".  for "result[ii + " = :  in ii ]result  result return |
| fake | | N/A | N/A | input :def)fun(  result [= ]  isalpha) result .ii[=(  input" in) split.for" ii(].  result return |
| fake | | N/A | N/A | input :def)fun(  result=result"][":  input) if (isalpha.not:  result(sorted=input+)  result return |
| fake | | N/A | N/A | input :def)fun(  result .is "digit= "  if )input(isdigit.:  result+ 2= result*  result return |
| fake | | N/A | N/A | input :def)fun(  input = result" " =  if!0"input".:=[ ]  upper) result .input=+(  result return |
| fake | | N/A | N/A | input :def)fun(  not = digit" result "  input) if (isdigit.not:  result+ 2= result*  result return |
| fake | | N/A | N/A | input :def)fun(  result."=. " .  split(input)if.1 "len)(:.">  result.split"input=1 ]("+) -.[  result return |
| fake | | N/A | N/A | input :def)fun(  result result" " =:  j=if"input[1] -":=  upper]result(input+1[ =.) -  result return |
| fake | | N/A | N/A | input :def)fun(  result "= i"  if )input(isalpha.:  upper) result .input=+(  result return |
| fake | | N/A | N/A | input :def)fun(  result result" " =:  a=if" 0"input:=[]  upper[ result( 0.input)]+=  result return |
| fake | | N/A | N/A | input :def)fun(  :result=result"]["  if )input(isalnum.:  result(sorted=input+)  result return |
| fake | | N/A | N/A | input :def)fun(  result>"= > " >  split(input)if.1 "len)(:.">  split( 0] input+result.".=)"[  result return |
| fake | | N/A | N/A | input :def)fun(  result =". "  input) if (isalpha.not:  input =result +  result return |
| fake | | N/A | N/A | input :def)fun(  result =". "  input +result* =(  isalpha)not.input()( )  result return |
| fake | | N/A | N/A | input :def)fun(  result "= i"  result + input=(  upper)isalpha)input.()*.(  result return |
| fake | | N/A | N/A | input :def)fun(  :result=result"]["  result+input=(  input)sorted)isalnum.*()(  result return |
| fake | | N/A | N/A | input :def)fun(  result>"= > " >  result(split.input=0 ].(+" [")  split(len.1)input)"(>"*)).(  result return |
| fake | | N/A | N/A | input :def)fun(  result result" " =:  0] result (input=+[  0)upper*a)input==("[.")](  result return |
| fake | | N/A | N/A | input :def)fun(  result .is "digit= "  result +result* =(  isdigit)2(input.*)  result return |
| fake | | N/A | N/A | input :def)fun(  input = result" " =  result + input=(  upper)0.input)*[=(!".")](  result return |
| fake | | N/A | N/A | input :def)fun(  not = digit" result "  result +result* =(  isdigit.2) input)not)(*(  result return |
| fake | | N/A | N/A | input :def)fun(  result."=. " .  input"1( result+split).- (.="][  split(len.1)input)"(>"*)).(  result return |
| fake | | N/A | N/A | input :def)fun(  result=result"][":  input*result=sorted((+)  isalpha)not.input()( )  result return |
| fake | | N/A | N/A | input :def)fun(  result result" " =:  1]result=input([ + -  input)upper(1)j=]-.")([=*"  result return |

Table S2. Activated clusters in each contrast

|  |  | peak MNI coordinates | | | Cluster size | | peak-p  (FWER) |
| --- | --- | --- | --- | --- | --- | --- | --- |
|  |  | X | Y | Z | vertices | mm2 |  |
| Real code > Fake code | |  |  |  |  |  |  |
| Left hemisphere | |  |  |  |  |  |  |
| Lateral prefrontal cortex (inferior/middle frontal gyri, precentral and superior frontal sulci) | | -51.7 | 16.8 | 21.1 | 1484 | 2962.45 | 0.0014 |
| Intra-parietal sulcus/supramarginal and angular gyri | | -45.4 | -54.1 | 39.9 | 846 | 1206.53 | 0.0084 |
| Middle/inferior temporal gyri and superior temporal sulcus | | -61.4 | -52.7 | -0.6 | 481 | 866.16 | 0.0128 |
| Calcarine sulcus/lingual gyrus | | -5.3 | -75.4 | 0.1 | 342 | 773.52 | 0.0150 |
| Right hemisphere | |  |  |  |  |  |  |
| Medial occipital cortex  Lingual gyrus/pericalcarine sulcus | | 13.2 | -89.9 | -0.4 | 904 | 2313.45 | 0.0036 |
| Angular gyrus | | 43.3 | -61.7 | 45.4 | 405 | 591.32 | 0.0356 |
| Inferior frontal sulcus  Peak: middle frontal gyrus | | 47.1 | 24.8 | 30.5 | 274 | 485.05 | 0.0448 |
| Language > Math | |  |  |  |  |  |  |
| Left hemisphere | |  |  |  |  |  |  |
| Superior temporal cortex | | -51.6 | -7.7 | -14.0 | 2854 | 5384.96 | 0.0002 |
| Medial occipital cortex  Peak: lingual gyrus | | -7.4 | -93.7 | -8.7 | 1606 | 3693.39 | 0.0002 |
| Pars triangularis | | -45.2 | 33.4 | -5.2 | 523 | 1070.54 | 0.0088 |
| Precuneus | | -4.7 | -49.5 | 27.8 | 445 | 811.74 | 0.0172 |
| Superior frontal gyrus | | -10.7 | 52.1 | 34.9 | 333 | 706.60 | 0.0224 |
| Right hemishpere | |  |  |  |  |  |  |
| Occiptial pole | | 12.8 | -92.0 | -7.1 | 932 | 2409.48 | 0.0006 |
| Anterior superior temporal sulcus | | 57.6 | -8.4 | -7.3 | 446 | 1073.59 | 0.0104 |
| Temporal pole | | 42.5 | 15.9 | -28.1 | 319 | 671.89 | 0.0222 |
| Posterior superior temporal sulcus | | 51.5 | -34.1 | 4.1 | 298 | 613.10 | 0.027 |
| Medial orbitofrontal sulcus | | 8.5 | 46.7 | -13.7 | 199 | 547.23 | 0.0336 |
| Precuneus | | 4.6 | -46.9 | 24.8 | 284 | 502.91 | 0.0376 |
| Math > Language | |  |  |  |  |  |  |
| Left hemisphere | |  |  |  |  |  |  |
| Intraparietal sulcus  Peak: supramarginal gyrus | | -50.4 | -38.9 | 44.0 | 1842 | 2797.80 | 0.0004 |
| Superior frontal sulcus | | -27.2 | 3.5 | 49.7 | 437 | 827.32 | 0.015 |
| Anterior middle frontal gyrus  Peak: inferior frontal sulcus | | -34.8 | 37.0 | 9.9 | 375 | 702.91 | 0.0204 |
| Superior insula sulcus | | -28.5 | 20.4 | 8.0 | 283 | 575.54 | 0.0308 |
| Inferior temporal gyrus | | -53.2 | -59.3 | -9.3 | 291 | 484.12 | 0.0408 |
| Right hemisphere | |  |  |  |  |  |  |
| Intraparietal sulcus | | 31.1 | -46.8 | 43.2 | 2502 | 3491.09 | 0.0002 |
| Anterior middle frontal gyrus | | 34.7 | 44.9 | 21.4 | 731 | 1406.92 | 0.0058 |
| Inferior temporal gyrus | | 45.5 | -60.4 | -8.2 | 591 | 1176.76 | 0.0088 |
| Middle anterior cingulate cortex  Peaks at the middle section of the pericallosal sulcus | | 5.4 | -1.6 | 30.2 | 502 | 1040.37 | 0.0092 |
| Superior frontal gyrus | | 21.1 | -4.0 | 59.1 | 504 | 934.90 | 0.011 |
| Anterior insula | | 35.5 | 16.8 | -1.2 | 351 | 688.23 | 0.0218 |
| Inferior precentral sulcus | | 48.4 | 5.8 | 27.7 | 257 | 549.18 | 0.0348 |
| Logic > Language | |  |  |  |  |  |  |
| Left hemisphere | |  |  |  |  |  |  |
| Intraparietal sulcus  Extends medially to precuneus and ventrally to inferior temporal gyrus | | -40.6 | -50.6 | 35.5 | 3347 | 5731.62 | 0.0004 |
| Pars orbitalis  Extends dorsally to precentral and middle frontal gyri | | -40.9 | 48.2 | -6.8 | 2120 | 4350.10 | 0.002 |
| Superior frontal gyrus | | -9.3 | 33.4 | 37.2 | 310 | 700.63 | 0.0334 |
| Pericallosal sulcus | | -7.4 | -32.0 | 30.0 | 333 | 640.21 | 0.0372 |
| Right hemisphere | |  |  |  |  |  |  |
| Intraparietal sulcus Extends medially to precuneus and ventrally to inferior temporal gyrus | | 31.0 | -47.0 | 40.2 | 3699 | 6446.9 | 0.0004 |
| Middle frontal gyrus Extends to superior frontal sulcus, precentral sulcus, and frontal pole | | 45 | 34.5 | 26.1 | 1908 | 3816.88 | 0.0036 |
| Pericallosal sulcus | | 8 | -36.3 | 29.1 | 343 | 683.83 | 0.0286 |
| MSIT | |  |  |  |  |  |  |
| Left hemisphere | |  |  |  |  |  |  |
| Fusiform gyrus  Extends to intraparietal sulcus | | -30.7 | -67.8 | -10.3 | 4059 | 7701.89 | 0.0002 |
| Superior precentral sulcus | | -22.1 | -0.9 | 47.0 | 421 | 891.27 | 0.0232 |
| Superior insula sulcus | | -33.8 | 23.1 | 11.1 | 339 | 690.26 | 0.0324 |
| Inferior precentral sulcus | | -49 | -2.5 | 37.8 | 314 | 605.36 | 0.0406 |
| Right hemisphere | |  |  |  |  |  |  |
| Intraparietal sulcus Extends to fusiform gyrus | | 40.9 | -40.5 | 35.8 | 3727 | 6865.08 | 0.0002 |
| Superior insula sulcus | | 29.7 | 26.8 | 4.3 | 438 | 838.39 | 0.0198 |
| Superior precentral sulcus | | 29.6 | -13.0 | 52.1 | 391 | 748.22 | 0.0232 |
| Middle anterior cingulate cortex Peak at medial superior frontal gyrus | | 10.5 | 12.0 | 47.7 | 313 | 598.73 | 0.0294 |
| Inferior precentral sulcus | | 43.1 | 2.4 | 29.8 | 231 | 484.89 | 0.043 |

Table S3. FDR-corrected p-values for the post-hoc paired t-tests among the overlap between code contrast and the localizer contrasts

| Contrasts  (Left hemisphere) | Language > math | Math > language | Logic > language |
| --- | --- | --- | --- |
| Math > language | 0.607 | -- | -- |
| Logic > language | 0.033 | 0.037 | -- |
| MSIT | 0.624 | 0.210 | 0.001 |

| Contrasts  (Right hemisphere) | Language > math | Math > language | Logic > language |
| --- | --- | --- | --- |
| Math > language | 0.895 | -- | -- |
| Logic > language | 0.103 | < 0.001 | -- |
| MSIT | 0.895 | 0.895 | 0.025 |
