## Supplementary figures and images for "Computer code comprehension shares neural resources with formal logical inference in the fronto-parietal network"

### supplementary figure S1

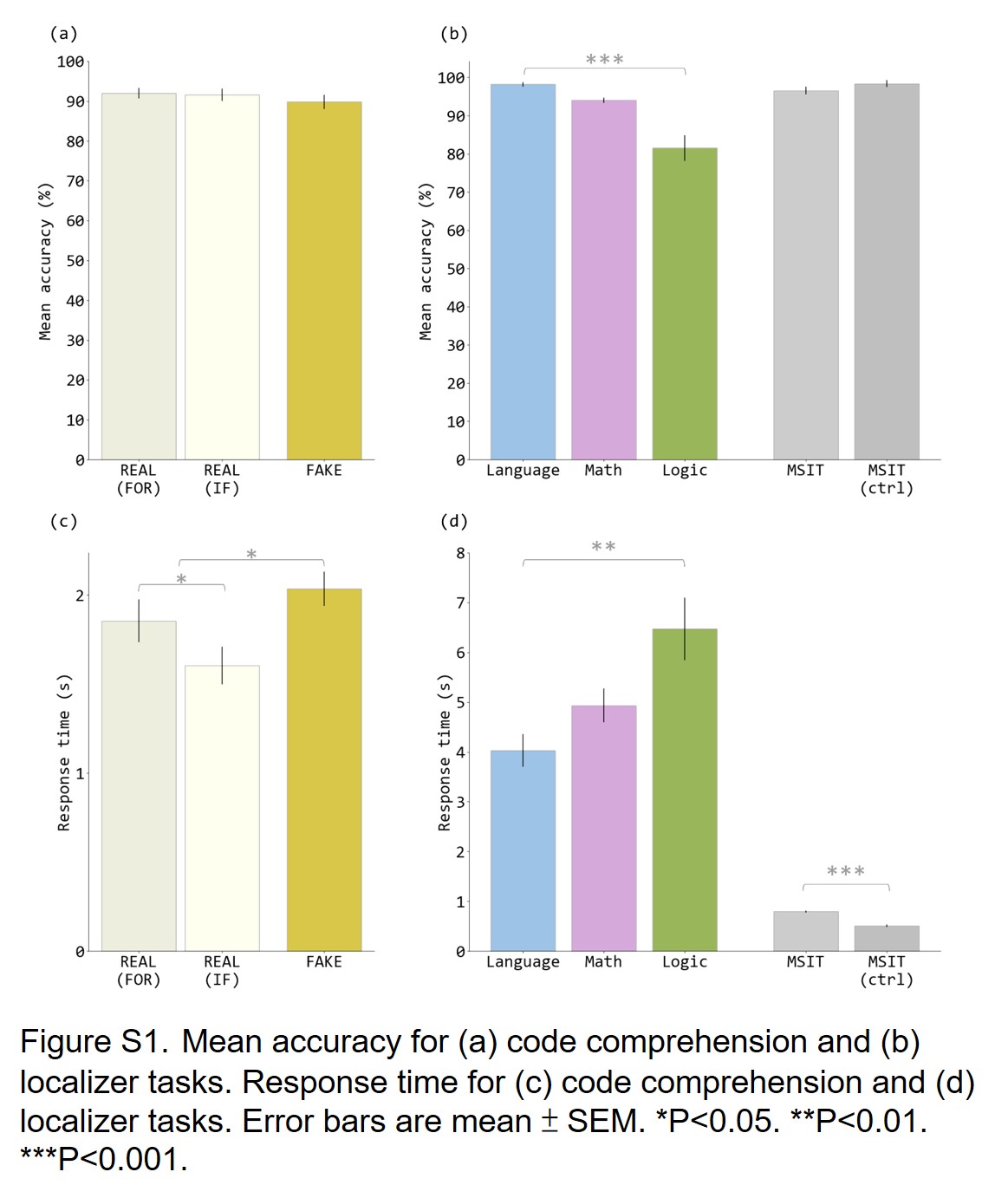
